# Natural Killer cell contractility and cytotoxicity is driven by TLR3 agonist-mediated TAZ cytoplasmic sequestration

**DOI:** 10.1101/2023.01.18.524654

**Authors:** Darren Chen Pei Wong, Zekun Xia, Nandi Shao, Jin Ye Yeo, Ivan Yow, T. Thivakar, Andres M. Salazar, Yih-Cherng Liou, Boon Chuan Low, Jeak Ling Ding

## Abstract

The evolutionarily conserved YAP/TAZ mechano-responsive transcription cofactors regulate development and tumorigenesis. Here, we show for the first time that human natural killer (NK) cells specifically express TAZ but not YAP, and TAZ serves to limit NK cytotoxicity. The TLR3 agonist, hiltonol, was able to trigger cytoplasmic sequestration of TAZ, which increased NK cytotoxicity. Mechanistically, a low dose of hiltonol enhanced the intracellular contractility of NK cells accompanied by an increase in active RhoA and myosin light chain phosphorylation, conceivably through an increase in ERK1/2 activation-dependent ROS production. Importantly, we showed that the dissociation of LATS1 from actin upon activation of contractility, is required to sequester TAZ in the cytoplasm. Functionally, hiltonol also reduced the NK cell surface inhibitory receptors, KIR3DL1 through c-Myc inhibition and PD-1 through enhanced contractility. Direct inhibition of c-Myc promoted NK cytotoxicity against K562 lymphoblast. We corroborated our findings by coculturing hiltonol-pretreated NK cells with breast and lung cancer cells, and demonstrated increased NK-mediated cancer killing, which conceivably occurred via sequestration of TAZ to the cytoplasm which facilitated NK cytotoxicity. Our findings pave the way for *ex vivo* rejuvenation of NK cells for *in vivo* immunotherapies, which could involve a cocktail of hiltonol and c-Myc inhibitor.

## 1. Introduction

The innate immune Natural Killer (NK) cells are indispensable in defending against infection and cancer^[1]–[4]^. Of particular interest is the integration of intrinsic and extrinsic cell and molecular, biochemical and biophysical signals, which activate dormant transcriptional networks responsible for NK cytotoxicity through various pattern recognition receptors (PRRs), for example, Toll-like receptors (TLRs) 1-9 in humans^[5]–[8]^. Upon activation, NK cells elicit their cytolytic capabilities mainly through the production of granzyme and perforin, and degranulation of these effectors to cytolyse target cells. Like other immune cells, NK cell activation has traditionally been associated with the integration of activating and inhibitory signals through several activating (e.g. NKG2D and DNAM-1) and inhibitory (e.g. PD-1, KIR family receptors and TIGIT) cell surface receptors^[2][9][10]^. However, several key questions remain on how the advanced stages of cancer cells evade these activating signals, through downregulation of NK cell cytotoxicity or promotion of NK cell exhaustion. Importantly, what are the extrinsic and intrinsic triggers that modulate NK cell activity to reinvigorate it for immunotherapy? We have recently identified intracellular contractility in NK cells which upregulates NK cytotoxicity by promoting nuclear localisation of transcription factor, Eomes^[1]^. Other studies also suggested that NK cells respond to mechanical forces^[11]^, indicating that increased substrate stiffness enhances NK cell cytotoxicity^[12][13]^. However, it is well known that cancer cells are softer than normal cells, especially for highly metastatic cancer cells^[14][15]^. Hence, it is of interest to delineate mechanosensitive pathways that could circumvent the reduced cancer cell stiffness and enhance NK cytotoxicity for potential therapeutic options.

Poly-ICLC (Hiltonol) is an advanced form of the synthetic poly-I:C dsRNA derivative stabilized with poly-lysine and carboxymethyl cellulose that effectively enters the endosome, thus making it a ‘reliable and authentic’ viral mimic that activates the human PRRs (e.g. TLR3 and MDA-5)^[16]^. The TLR3 in NK cells was first identified to recognise dsRNA, a viral replication intermediate, and mediates pro-inflammatory responses through type I interferons (IFNs) and chemokines^[8][17][18]^. However, recent studies suggest that upon stimulation by dsRNA, the activated TLR3 regulates more than just the production of IFNs and chemokines. In fact, activation of TLR3 on NK cells can induce dendritic cell maturation, which is consistent with *Tlr3* null mice being hypo-responsive to cytokine stimulation and unable to limit cancer metastasis^[8]^. These observations suggest that activation of NK cell TLR3 induces NK-cancer killing activity. Several clinical trials using TLR3 agonists as an adjuvant for therapeutic vaccination against cancers are underway^[19][20]^. Despite extensive studies on the role of TLR3 in inducing immune responses through inflammation, the mechanisms underlying TLR3-induced NK cytotoxicity has remained unclear. Interestingly several conflicting studies implicate the role of TLRs in regulating cellular contractility in aortic vascular smooth muscle cells and cardiomyocytes^[21][22]^. To seek clarity and to fill the knowledge gap, a study of contractility and mechanosensitive pathways regulated by NK contractility is pertinent and essential. Moreover, the important, yet variegated roles of the cytoskeleton in NK cell are increasingly gaining attention^[11][23]–[26]^.

Various NK cell immunodeficiency diseases are purportedly caused by the mis-regulation of cytoskeletal proteins^[27]^. We have previously shown that despite lacking focal adhesions, NK cells can remarkably alter intracellular contractility for nuclear localisation of the transcription factor, Eomes, and elicit early cytotoxic responses against cancer cells^[1]^. Furthermore, biochemical and biophysical signals should be integrated to regulate pathophysiological activities. For instance, the Hippo-Yes-associated protein (YAP)/WW domain-containing transcription regulator 1 (WWTR1/ TAZ) pathway is biochemically and biophysically co-regulated^[28]–[30]^.

The Hippo-YAP/TAZ pathway was first identified as a growth regulator and its dysregulation was reported to promote tumorigenesis^[29][31]^. YAP and TAZ are transcription cofactors that are canonically inhibited by the Hippo pathway through a series of kinases (MST and LATS) which are sequentially activated by phosphorylation^[30][32]^. Recent findings suggest that YAP and TAZ are mechano-responsive, and their intracellular compartmentalisation can be affected by mechanical forces^[33]^. However, the mechanistic roles of Hippo-YAP/TAZ in immune cells have hitherto remained unclear. Recent evidence suggests that YAP suppresses CD8 T-cell differentiation and function^[34]^. However, how and whether Hippo-YAP/TAZ affects NK cell activity, needs clarification. Moreover, the triggers and mechanisms that activate or inactivate the Hippo pathway to regulate YAP/TAZ activity in NK cells are unknown. Since TLR3 activation was shown to induce endothelial cell contractility^[22]^ and YAP/TAZ are mechano-regulated^[29][35]^, it is possible that NK cell contractility can control YAP/TAZ through mechanotransduction events. This prompted us to hypothesise that TLR3 activation regulates YAP/TAZ subcellular localisation through enhanced NK cell contractility, which potentially modulates NK cytotoxicity.

Here, using primary human NK (hNK) cells and an NK cell line undergoing clinical trial (NK-92), we show that NK cells uniquely express TAZ but not YAP mRNA and protein. Overexpression and knockdown of TAZ suppressed and enhanced NK cell cytotoxicity, respectively, suggesting that inhibition of TAZ could heighten NK cytotoxicity. Next, we showed that treatment of NK cells with TLR3 agonist, hiltonol, promoted the cytoplasmic compartmentalisation of TAZ, which was associated with increased NK cytotoxicity. Mechanistically, we used multi-modal approaches to provide evidence that hiltonol activates ERK1/2 to enhance intracellular contractility. This was demonstrated by an increase in F-actin tensional force in laser ablation experiments and an increase in RhoA-MLC2 activation. Our results showed that: (i) activated ERK1/2 raised the mitochondrial ROS level; (ii) enhanced NK contractility released LATS1 kinase from actin which correlated with LATS1 phosphorylation at Ser89; and (iii) phosphorylated LATS1 then phosphorylated and inhibited TAZ which was sequestered in the cytoplasm. We also identified c-Myc, a known downstream target of TAZ activation, to conversely limit NK cytotoxicity and found that in the presence of nuclear TAZ, c-Myc activity maintained a high level of inhibitory receptor, KIR3DL1, on NK cells. On the other hand, inhibition of c-Myc directly or through hiltonol treatment, reduced NK inhibitory receptor to unleash the cytotoxic potential of NK cells against cancer cells. Taken together, we have demonstrated, in a lymphoblast cell line and two cancer cell types (breast and lung), that hiltonol treatment increased NK cell cytotoxicity against cancer cells. Our findings provide potentials for immunomodulating NK cells to kill cancers.

## 2. Results

### 2.1 TAZ limits NK cell cytotoxicity

Since YAP and TAZ are homologs with similar structure and function^[28][29]^, we first verified if YAP and TAZ are equally expressed in NK cells. Surprisingly, we could only detect TAZ but not YAP expression at both the mRNA (**Figure 1A**) and protein (**Figure 1B**) levels in both NK-92 and primary human NK (hNK) cells. Upon cross referencing with the available online database^[36]^, we confirmed that indeed NK cells only expressed TAZ but not YAP (**Figure S1**).

**Figure 1.**
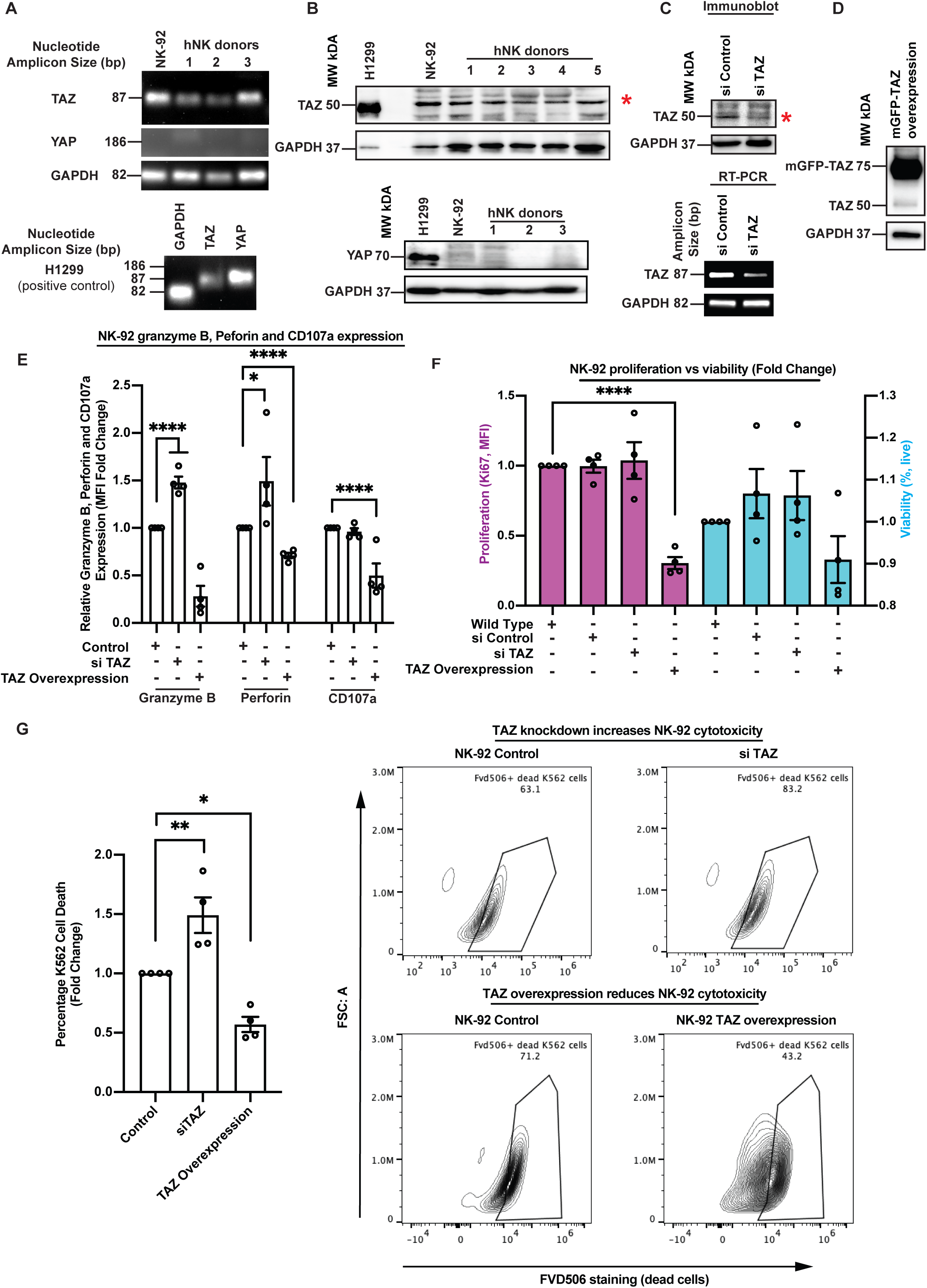
TAZ limits NK cell cytotoxicity. **(A)** RT-PCR analysis shows that NK-92 cell line and three primary human NK (hNK) cells from healthy donors express TAZ but not YAP. The H1299 NSCLC cells (positive control) express both TAZ and YAP. All RT-PCR analyses were repeated thrice, and a representative gel blot was shown. The size markers represent the nucleotide amplicon size and not gene size. **(B)** Immunoblot shows NK-92 and hNK cells express TAZ protein but not YAP protein. H1299 NSCLC was used as a positive control. All blots were repeated thrice, and a representative image was shown. **(C)** Immunoblot and RT-PCR analyses show knockdown of TAZ protein and mRNA levels in NK-92 cells treated with TAZ siRNA. The red asterisk indicates the TAZ band. The size markers for RT-PCR images represent the nucleotide amplicon size and not gene size. **(D)** Immunoblot shows TAZ overexpression in NK-92 cells. **(E)** Taz knockdown increased granzyme and perforin expression in NK-92 cell, while TAZ overexpression reduced expression of granzyme, perforin and surface CD107a. One-way ANOVA was used to compare the different conditions. The graph represents means ± SEM and n=4 for all conditions. **(F)** The proliferation (purple) of NK-92 cells, as indicated by Ki67, decreased only with TAZ overexpression. However, the viability (blue) of NK-92 cells were not affected after TAZ knockdown or overexpression. One-way ANOVA was used to compare the different conditions. The graph represents means ± SEM and n=4 for all conditions. **(G)** (Left) TAZ knockdown increased NK-92 cell cytotoxicity (higher K562 death) while TAZ overexpression reduced NK-92 cell cytotoxicity. For TAZ overexpression, K562 target cells were stained with CFSE-Far Red. One-way ANOVA was used to compare the different conditions. The graph represents means ± SEM and n=4 for all conditions. (Right) Representative flow cytometric graphs show TAZ knockdown and TAZ overexpression increased and decreased NK-92 cytotoxicity, respectively. Refer to Figure S2 for cytotoxicity assay gating strategy using FVD506 dead cell staining. For all graphs, *p < 0.05, **p < 0.01 and ****p < 0.0001.

Being homologous, YAP and TAZ are apparently regulated by similar mechanisms in various systems^[29][30]^. Since YAP has been reported to elicit a broad inhibitory role in T cells (CD4 and CD8)^32-34^, which are of the common lymphoid lineage including innate NK cells, we hypothesised that TAZ might also regulate NK cytotoxicity. To test this, we knocked down (**Figure 1C**) and over-expressed TAZ (**Figure 1D**) in NK-92 cells. We observed significant up- and down-regulation of key cytotoxic molecules (granzyme b and perforin) and the surface degranulation marker CD107a in TAZ knockdown and overexpression in resting NK-92 cells, respectively (**Figure 1E**). Interestingly, the proliferation rate of NK-92 cells, as indicated by Ki67, remained unchanged in either wild type or siRNA control or TAZ knockdown NK-92 cells but decreased significantly in TAZ-overexpressing cells (**Figure 1F**). Nevertheless, the viability of NK-92 cells did not change with either TAZ knockdown or overexpression. To ascertain the functionality of knockdown /overexpression of TAZ in NK-92 cells, we performed a killing assay using the Fixable Viability Dye eFluor™ 506 dye that irreversibly labels dead cells (Fvd506 positive, see **Figure S2** for gating strategy) and can be fixed and washed to stain for intracellular antigens without any loss of staining intensity of the dead cells^[1][37]^. The myelogenous leukaemia K562 model cell line was stained with Fvd506 dye after coculturing with NK-92 cells for four hours and NK-92 TAZ-knocked down cells demonstrated enhanced killing efficiency against K562 cells (**Figure 1G; middle bar**). Conversely, overexpression of TAZ reduced the killing efficiency of NK-92 cells on K562 cells (**Figure 1G; right bar)**.

Together, our results showed that NK cells express TAZ but not YAP and that overexpression of TAZ limits the cytotoxic potential of NK cells. Since TAZ is a transcriptional co-activator, we reasoned that understanding the conditions that influence TAZ modulation, for example its cellular compartmentalisation would be mechanistically enlightening. Henceforth, we focused on possible factors altering TAZ subcellular localization in NK cells.

### 2.2 TLR3 activation by hiltonol induces cytoplasmic compartmentalisation of TAZ and increases NK cytotoxicity

Although first identified to be an anti-viral receptor responding to viral dsRNA^[8][20]^ recent studies suggest that TLR3 activation in NK cells elicit cytotoxic effects on cancer^[8][17][38]^. Other studies also demonstrated that TLR3 activates cellular contractility in non-immune systems^[18][21][22]^. Since YAP/TAZ are mechano-sensitive and TLR3 has potential anti-tumour capabilities, we reasoned that activation of TLR3 might regulate the subcellular localization of TAZ.

To test whether TLR3 activation affects TAZ subcellular localization, we treated NK cells with increasing doses of hiltonol. We observed a significant dose-dependent increase in TLR3 expression in hiltonol-treated hNK cells (**Figure S3A**). Confocal images of TAZ localization showed that hiltonol induced a significant cytoplasmic sequestration of TAZ, corresponding to the dose of hiltonol, in both NK-92 (**Figure 2A**) and hNK cells (**Figure 2B**), but without affecting cell viability (**Figure S3B**). Interestingly, in control resting NK cells, TAZ was homogeneously present in both the cytoplasm and nucleus of NK cells, in contrast to being predominantly present in the nucleus of other adherent cell types^[39][40]^. To corroborate our observations, we treated NK-92 and hNK cells with hiltonol and performed imaging flow analysis to empirically quantify the overlap signal of nuclear stain (DAPI) to TAZ (**Figure 2C-F**). We quantified lesser overlap in nuclear signal of TAZ (represented by median similarity TAZ/DAPI score) in NK cells treated with hiltonol, at a level which was consistent with the cytoplasmic localisation of TAZ. To confirm that the compartmentalization of TAZ to the cytoplasm is not due to an increase in nuclear size, we quantified the nuclear size of hNK cells treated with hiltonol. Interestingly, the nucleus was significantly smaller in hiltonol treated hNK cells (**Figure S3C, D**), thus authenticating the ratio of TAZ distribution between nucleus and cytoplasm. Next, we performed a western blot analysis of TAZ and LATS1 (the kinase that inactivates TAZ) and showed increased phosphorylation of both LATS1 and TAZ (**Figure 2G**). This implies activation of LATS1 (phosphorylation at Serine 909) and inhibition of TAZ (phosphorylation at Serine 89) in NK cells treated with hiltonol.

**Figure 2.**
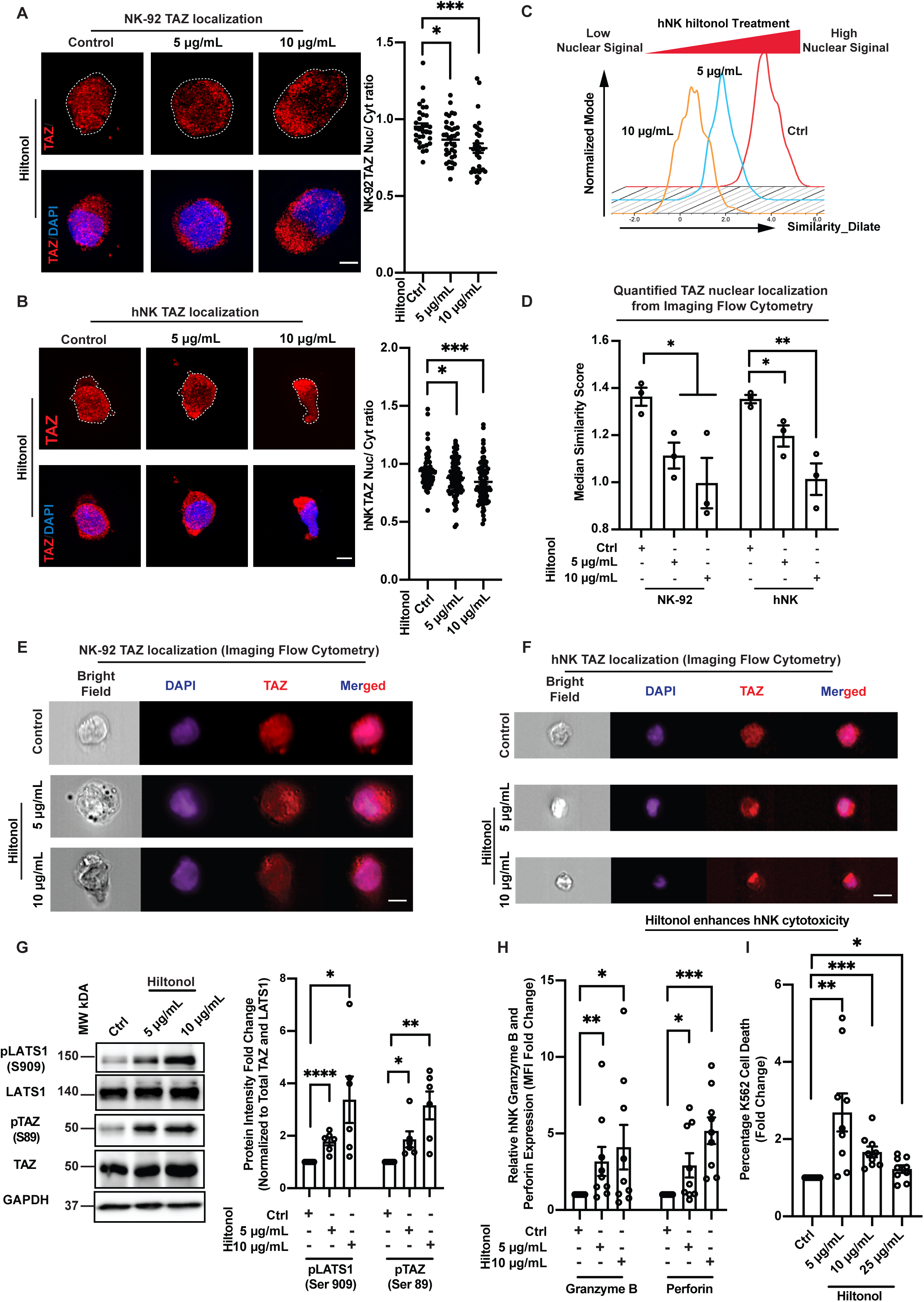
TLR3 activation by hiltonol induces cytoplasmic compartmentalization of TAZ and increases NK cytotoxicity. **(A)** Immunofluorescence staining of TAZ in NK-92 cells shows increased cytoplasmic localization of TAZ with hiltonol treatment, Scale bar = 5 μm. The white dotted lines demarcate cell periphery. Quantification on the right represents means ± SEM and n = 32, 36 and 31 for ctrl (control), 5 μg/mL hiltonol and 10 μg/mL hiltonol, respectively. **(B)** Immunofluorescence staining of TAZ in hNK cells shows increased cytoplasmic localization of TAZ upon hiltonol treatment. Scale bar = 5 μm. The white dotted lines demarcate cell periphery. Quantification on the right represents means ± SEM and n = 74, 82 and 85 for ctrl, 5 μg/mL hiltonol and 10 μg/mL hiltonol, respectively. Data are representative of three hNK donors. **(C) and (D)** Representative imaging flow cytometric histogram (C) and quantification (D) of E and F, showing TAZ compartmentalisation to the cytoplasm of NK-92 (E) and hNK (F) cells with hiltonol treatment. The graph represents means ± SEM, and n=3 for NK-92 cells, and three hNK donors were used for hNK cells. For each n, 3000 to 5000 cells were imaged for quantification. The “similarity dilate” values were calculated according to nuclear translocation wizard in IDEAS 6.0 software. **(E) and (F)** Representative images of imaging flow cytometry showing TAZ compartmentalisation to the cytoplasm of NK-92 (E) and hNK (D) cells with hiltonol treatment. Scale bar = 10 μm. **(G)** Immunoblot analysis shows increased LATS activation (pLATS1) and increased TAZ inactivation as pTAZ increases in NK-92 cells treated with hiltonol. The graph represents means ± SEM, and n=6. Student’s T-test was used to compare different conditions to control. **(H)** Hiltonol treatment increases hNK granzyme B and perforin expression as indicated by expression fold-change of the MFI (mean fluorescence index). The graph represents means ± SEM, and n=9 individual experiments representative of three hNK donors. **(I)** Hiltonol treatment (from 5 to 25 μg/mL) increases then decreases hNK cytotoxicity towards model target, K562 lymphoblast cells. The graph represents means ± SEM, and n=9 individual experiments representative of three hNK donors. For all graphs, *p < 0.05, **p < 0.01. ***p<0.001 and ****p < 0.0001 and One-way ANOVA was used to compare the different conditions unless otherwise stated.

Since knockdown of TAZ increases NK cytotoxicity, we reasoned that hiltonol-induced TAZ cytoplasmic compartmentalisation should correspond to a reduction in TAZ-mediated nuclear activity resulting in enhancement of NK cytotoxicity. Simultaneously, we showed that hiltonol treatment induced significant nuclear localization of Eomes (**Figure S3E**), a transcription factor responsible for reduced PD-1 expression^[3]^ which resulted in early enhancement of NK cytotoxicity against cancer^[1]^. To determine the functional effect of hiltonol on NK cells, we pre-treated NK cells with hiltonol and performed a cytotoxicity assay with K562 cells. There was a significant increase in the: (a) production of granzyme and perforin, (b) killing of K562 cells, and (c) NK-K562 binding (**Figure 2H, I and S3F, G**). Interestingly, a high dose of hiltonol (25 μg/mL) reversed its enhancement of NK cytotoxicity (**Figure 2I**). This was consistent with reports that high dosage of Poly-I:C/ hiltonol tends to reduce the cytotoxicity against target cells^[41]–[43]^. Henceforth, we focused on 5 and 10 μg/mL of hiltonol for experiments on functional analyses (e.g. cancer killing assays).

Altogether, we showed that TLR3 activation by hiltonol resulted in significant induction of cytoplasmic compartmentalisation of TAZ, which was associated with enhanced NK cytotoxicity. Next, we aimed to determine mechanistically, how TLR3 activation induced TAZ cytoplasmic compartmentalisation.

### 2.3 Hiltonol induces NK cell contractility and ROS production through ERK1/2

The study of TLR3 signalling in immune cells has thus far mainly focused on its downstream pro-inflammatory effects^[7][8][44]^. However, the mechanisms associated with TLR3 signalling, for example on cellular contractility, has hitherto remained ambiguous, although reported^[18][21][22]^. We have previously demonstrated that the presence of cancer cells enhanced intracellular contractility of NK cells which promoted nuclear compartmentalization of Eomes^[1]^. Here, our observation that hiltonol induces Eomes nuclear localization (**Figure S3E**) hints that hiltonol treatment activated intracellular contractility for early cytotoxicity response. An enhancement of intracellular contractility was also indicated by the observation that hiltonol treated NK cells have a smaller nuclear area (**Figure S3C, D**) and this is supported by a previous study which suggested that intracellular contractility compresses the nucleus^[45]^. TAZ is a transcription cofactor that can also be mechanically regulated^[28][29]^; thus we hypothesised that hiltonol treatment simultaneously induces a concerted effect on a biochemical pathway (through LATS1 phosphorylation) and a mechanical pathway (through enhancement of intracellular contractility).

To verify whether and how contractility was enhanced, we treated NK-92 cells with hiltonol and assayed for actin recoil velocity by performing laser ablation on actin filaments (**Figure 3A**). The recoil velocity should directly correlate with intracellular tensional force/contractility exerted by actin filaments^[1][46]^. We found that hiltonol dose-dependently induced a significant increase in the NK-92 actin recoil velocity (**Figure 3B**). Likewise, hNK cells showed a similar dose-dependent response (**Figure 3C**). Since the non-muscle myosin light 2 (MLC2) chain generates intracellular contractile forces through RhoA activation, we next verified if MLC phosphorylation and RhoA activation (RhoA-GTP) were induced by hiltonol treatment. We found that hiltonol significantly increased both the MLC2 phosphorylation (**Figure 3D**) and the active form of RhoA (RhoA-GTP) (**Figure 3E**).

**Figure 3.**
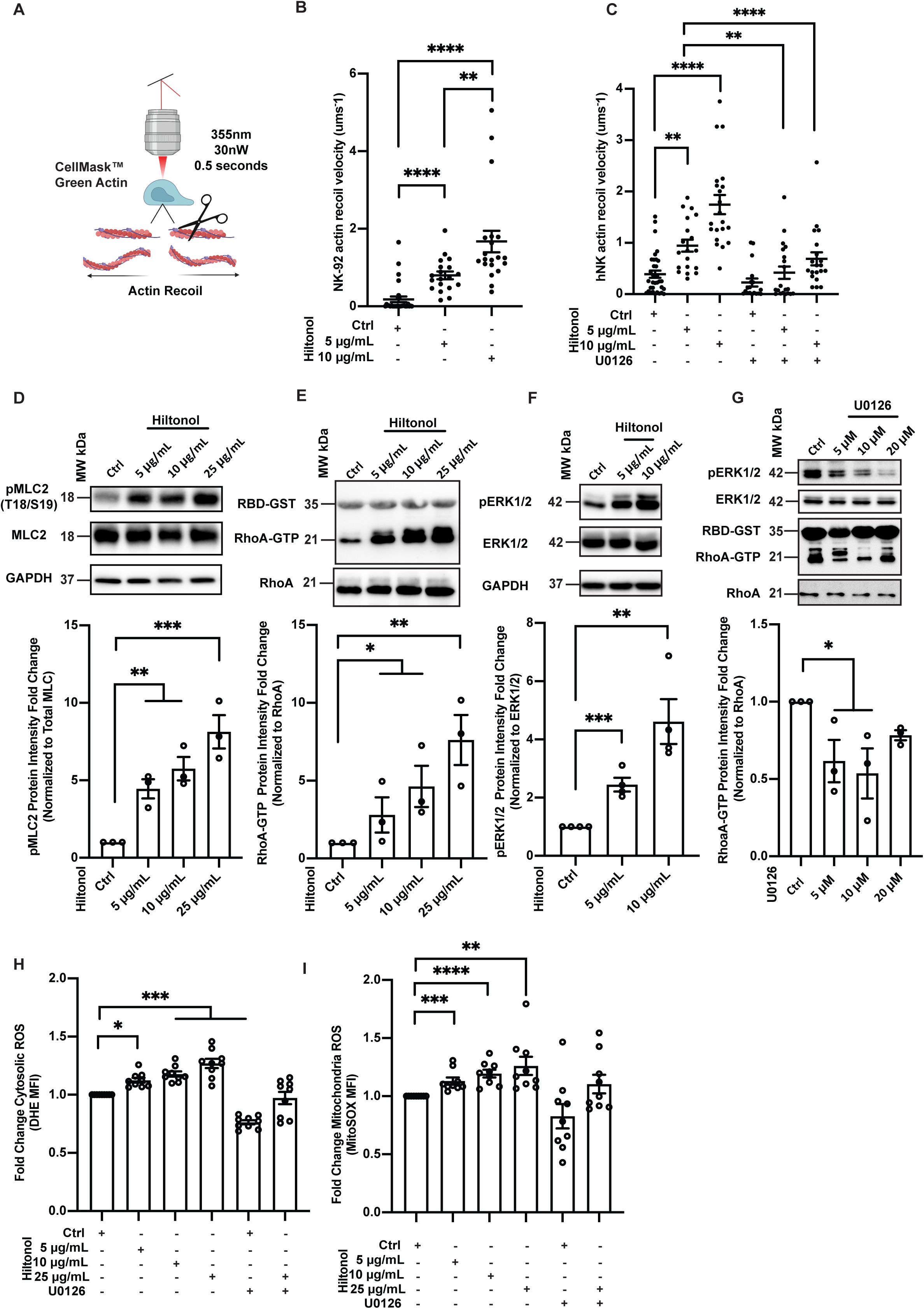
Hiltonol induces NK cell contractility and ROS production through ERK1/2. **(A)** Schematic showing laser ablation of NK cells with actin filaments labelled with CellMask™ green actin. The scissors represent laser ablation site and arrows represent direction of actin recoil. Details can be found in the materials and methods section “Laser ablation of NK cell actin filaments”. **(B)** Hiltonol treatment increases the initial recoil velocity of laser ablated NK-92 actin. The graph represents means ± SEM, and n=29, 19, 20 for ctrl, 5 μg/mL hiltonol and 10 μg/mL hiltonol, respectively. **(C)** Hiltonol treatment increases the initial recoil velocity of laser ablated hNK actin. ERK inhibitor U0126 (10 μM) reduced the increase in initial recoil velocity. The graph represents means ± SEM, and n=34, 19, 21, 18, 20, 20 for each column of the graph, respectively. **(D-F)** Immunoblots shows that hiltonol treatment increases NK-92 pMLC2 (phosphorylated myosin-like chain 2) that is correlated with RhoA activation (RhoA-GTP) and ERK activation (pERK1/2). The graph below each immunoblot represents means ± SEM, and n ≥3. Students T-test was used to compare different conditions to control. (**G**) Immunoblot showing that U0126 treatment reduces pERK1/2 levels, which is correlated to reduced RhoA activation (RhoA-GTP). The graph below represents means ± SEM, and n=3. Students T-test was used to compare different conditions to control. **(H and I)** Hiltonol treatment increases hNK cytosolic ROS as indicated by DHE staining (H) and mitochondria ROS as indicated by MitoSOX staining (I). This was inhibited with the ERK inhibitor U0125. The graph represents means ± SEM, and n=9 representative of three hNK donors. For all graphs, *p < 0.05, **p < 0.01. ***p<0.001 and ****p < 0.0001 and One-way ANOVA was used to compare the different conditions unless otherwise stated.

Next, we determined the mechanism by which TLR3 signalling activates intracellular contractility. It has been reported that systemic activation of mouse TLR3 resulted in ERK1/2 activation and MLC phosphorylation in aortic vascular smooth muscle cells^[22]^. We perceived that this could be conserved in NK cells since the common lymphoid progenitor from which NK cells differentiated from, are derived from the mesoderm, like muscle cells. To verify this possibility, we analysed ERK1/2 phosphorylation after hiltonol treatment, showing a dose-dependent increase in ERK1/2 phosphorylation (**Figure 3F**). Next, we directly inhibited ERK1/2 phosphorylation with U0126 and measured the actin recoil velocity. Inhibition of ERK1/2 with U0126 drastically reduced hiltonol-induced intracellular contractility (**Figure 3C**). Furthermore, treatment of NK-92 cells with U0126 (5 and 10 μM) significantly reduced RhoA activation, showing lesser RhoA-GTP (**Figure 3G**). Hence, ERK1/2 activation is responsible for RhoA-induced intracellular contractility in NK cells.

ROS production has been shown to activate ERK1/2^[47][48]^. However, recent evidence suggest that ERK1/2 could also induce ROS production^[49][50]^, indicating a positive feedback loop. A moderate increase in the level of ROS in NK cells was also proposed to improve NK cell division and function^[51]–[53]^. Hence, whether ERK1/2 activation directly correlates with ROS production after hiltonol treatment is of potential interest. We treated hNK cells with hiltonol and/ or U0126, and measured the cytoplasmic and mitochondrial ROS production with Dihydroethidium (DHE) staining and MitoSOX staining, respectively. Interestingly, both DHE and MitoSOX staining showed an increase in cytoplasmic ROS production, which was inhibited by U0126 (**Figure 3H, I**). These observations support our finding that hiltonol directly enhances NK cytotoxicity (Figure 2H, I). Nevertheless, the ROS increase continued to be significant at 25 µg/mL of hiltonol treatment (Figure 3H, I), although higher dose of hiltonol reduced NK cytotoxicity (Figure 2I). Hence, high ROS may limit NK activation/ cytotoxicity.

Taken together, we have shown that activation of TLR3 by hiltonol induced intracellular contractility through ERK1/2, with possible synergy between ERK1/2 activation and ROS production, which corresponded to the increase in NK cell cytotoxicity. Next, it was pertinent to elucidate how enhanced contractility promoted LATS1 phosphorylation, which in turn affected TAZ intracellular localisation.

### 2.4 Activation of intracellular contractility by hiltonol releases LATS1 from actin and sequesters TAZ in the cytoplasm

It was previously shown that the TAZ homolog, YAP, responds to mechanical forces and compartmentalises to the nucleus of fibroblast cells^[33]^. However, we have thus far, demonstrated that enhanced contractility (Figure 3A-E) correlated with TAZ cytoplasmic compartmentalisation, and promoted LATS1 phosphorylation (Figure 2). We reasoned that intracellular contractility in NK cells probably activated a concerted mechanism linking mechanical contractility to the biochemical regulation of TAZ.

Since ERK1/2 activation was associated with hiltonol-induced RhoA activation and contractility, we first inhibited ERK1/2 with U0126 and analysed TAZ and LATS1 phosphorylation. As shown in **Figure 4A**, U0126 treatment reduced TAZ and LATS1 phosphorylation in NK-92 cells, suggesting an increase in TAZ nuclear localization. Next, we tested whether direct pharmacological inhibition of myosin-based contractility with blebbistatin would impact the subcellular localization of TAZ in hNK cells. Indeed, we observed a significant nuclear re-compartmentalisation of TAZ when myosin-contractility was inhibited by blebbistatin (**Figure 4B**). This is in contrast to hiltonol induced TAZ cytoplasmic localization as demonstrated in Figure 2. Furthermore, Western blot analyses showed that hiltonol-induced LATS1 phosphorylation was also inhibited by blebbistatin (**Figure 4C**). These observations suggest an association between cell force (intracellular contractility) and promotion of TAZ sequestration in the cytoplasm, which is in line with a previous study showing blebbistatin treatment-induced nuclear localization of YAP/TAZ in MDCK cells^[54]^.

**Figure 4.**
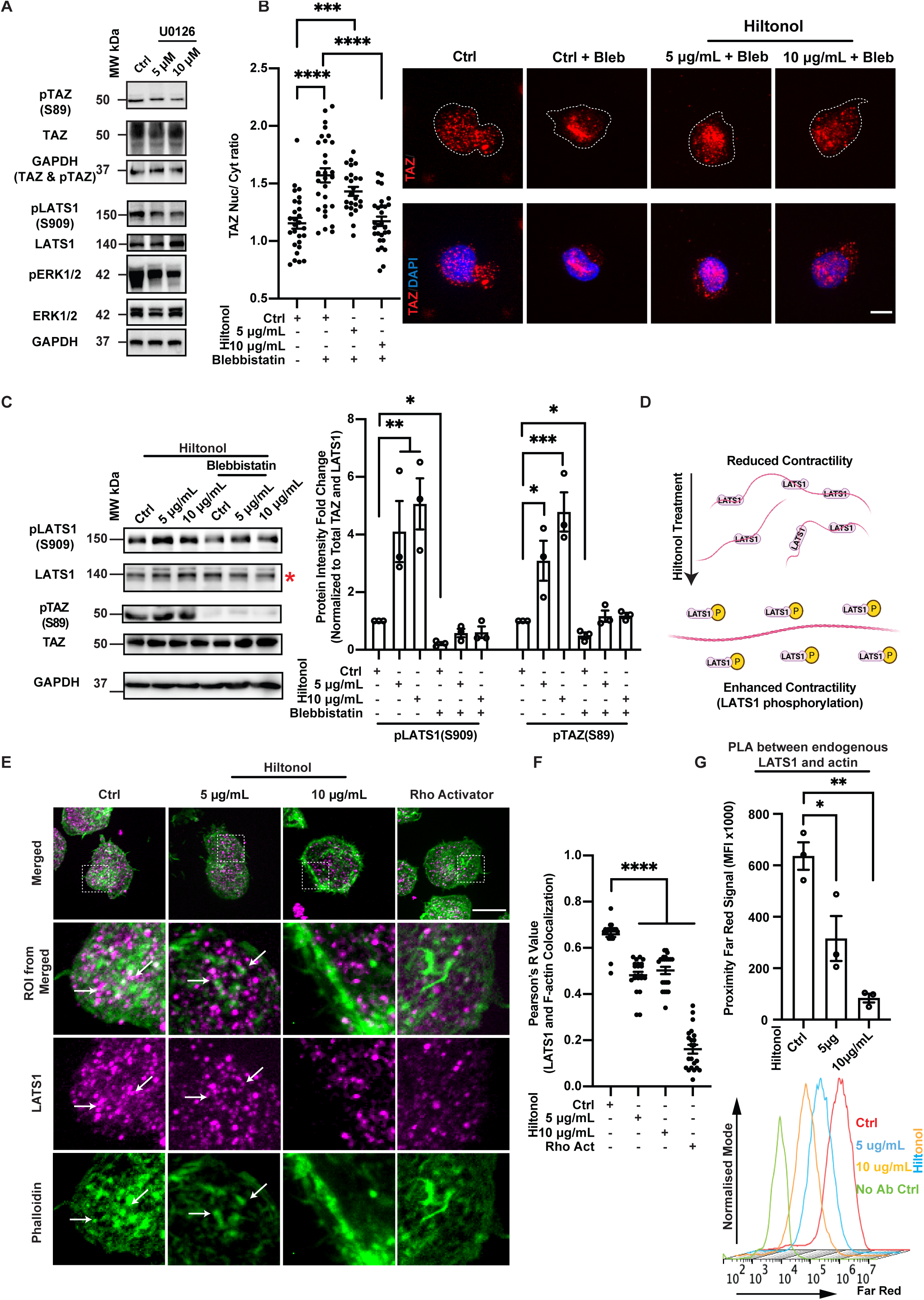
Hiltonol activation of intracellular contractility releases LATS1 from actin and sequesters TAZ in cytoplasm. **(A)** Immunoblot showing U0126 treatment (10 μM) overnight reduces TAZ phosphorylation and LATS1 phosphorylation in NK-92 cells, n= 3. **(B)** Quantification (Left) and representative images (Right) shows that myosin inhibitor, blebbistatin (40 μM) treatment overnight, increases TAZ nuclear localization in NK-92 cells. The white dotted lines outline the cell periphery. Scale bar = 5 μm. The graph represents means ± SEM, and n=27, 29, 24, 29 for each column. **(C)** Immunoblot shows that increases in pLATS1 and pTAZ with hiltonol treatment were blocked by the addition of blebbistatin (40 μM). The red asterisk indicates the LATS1 protein band. The graph represents means ± SEM and n=3. Students T-test was used to compare different conditions to control**. (D)** Schematic showing enhanced contractility (e.g. with hiltonol treatment) releases LATS1 from actin and correlates with increased LATS1 phosphorylation. **(E) and (F)** Representative immunofluorescence images of NK-92 cells (D) and quantification of Pearson’s R value on the right (E) shows reduced LATS1 and F-actin (phalloidin) colocalization with hiltonol treatment. Rho activator (1.5 μg/mL) was used as a positive control. Scale bar = 10 μm. The dotted white box represents the ROIs (regions of interest), which are magnified in panels below. The white arrows highlight points of colocalization. The graph represents means ± SEM and n= 22 for all columns. **(G)** Hiltonol treatment of NK-92 reduces PLA (proximity ligation assay) far red MFI signals, indicating reduced colocalization of LATS1 and actin. The graph represents means ± SEM and n= 3. The representative flow cytometry histogram below shows left shift (lesser signal) of peaks with hiltonol treatment. For all graphs, *p < 0.05, **p < 0.01. ***p<0.001 and ****p < 0.0001 and One-way ANOVA was used to compare the different conditions.

LATS1 kinase is an actin-binding protein which antagonises actin polymerisation^[55]^, suggesting that LATS1 activation through phosphorylation requires either (i) actin depolymerisation or (ii) its release from F-actin filaments. Since LATS1 is a direct inactivation kinase of TAZ but was activated with hiltonol treatment (Figure 2G), it is conceivable that enhanced contractility would ‘release’ LATS1 from actin and the freed LATS1 becomes phosphorylated (**Figure 4D**). Furthermore, studies on GPCRs identified the integrity of the actin cytoskeleton, a repressor of the Hippo pathway^[56]^. To test this hypothesis, we performed spinning disk confocal imaging analysis and calculated the Pearson’s R-Value between LATS1 and F-actin using Rho Activator as a positive control of increased contractility^[1]^. Indeed, treatment of NK cells with hiltonol or Rho activator (CN03) significantly reduced the correlation between LATS1 and F-actin (**Figure 4E, F**). To corroborate these observations, we performed a flow cytometric proximity ligation assay (PLA). PLA combined with flow cytometry allows quantitative analysis of proteins in proximity, as close as 40 nm^[57]^. We found that treating NK cells with hiltonol significantly reduced the association of LATS1 and actin (**Figure 4G**), implying that enhanced intracellular contractility releases LATS1 from F-actin.

Thus far, we have shown that NK cells express TAZ, and hiltonol treatment sequesters TAZ to the cytoplasm. Activation of contractility through ERK1/2 resulted in LATS1 dissociation from actin, which correlated with higher LATS1 phosphorylation and concomitant TAZ phosphorylation. To recapitulate, enhanced contractility induced by hiltonol treatment increased NK cell cytotoxicity. We next sought to determine how TAZ limits NK cell cytotoxicity.

### 2.5 TAZ-induced c-Myc activity regulates NK cell inhibitory receptor, KIR3DL1

One of the mechanisms by which NK cytotoxicity is regulated involves the balance of activating (e.g. NKG2D, DNAM-1) and inhibitory (e.g. NKG2A, TIGIT, PD-1, KIR2DL1 and KIR3DL1) surface receptors ^[2][10]^. We hypothesised that hiltonol-induced TAZ cytoplasmic sequestration could regulate the expression of one or multiple surface receptors on NK cells. To test this, we analysed the expression of various major surface receptors. Interestingly, while no upregulation of NK activating receptors was observed (**Figure 5A**), we noticed significant downregulation of the inhibitory receptors like PD-1 and KIR3DL1 (**Figure 5B**). In addition, the downregulation of PD-1 was consistent with Eomes-mediated suppression of PD-1^[3]^, and here we have also shown that Eomes localises to the nucleus of hiltonol-treated NK cells (**Figure S2C**), which was associated with the suppression of PD-1. PD-1 has been reported to be expressed on primary human NK cells^[58]^, making it an invaluable target for immunotherapy. In addition, to reconcile these observations with NK functional outcome, we analysed CD56 (NK maturation marker) and CD16 (NK antibody-dependent cell-mediated cytotoxicity marker) expressions on NK cells and showed that hiltonol treatment did not alter their expressions (**Figure S4**). On the other hand, the downregulation of KIR3DL1 raised the possibility that a transcription cofactor, TAZ, might have induced downstream genes that regulated the expression of KIR3DL1. We further confirmed this by knocking down TAZ in NK-92, which significantly reduced the surface expression of KIR3DL1 (**Figure 5C**). To strengthen our claim, we inhibited TAZ activity with the clinical photosensitiser and YAP/TAZ inhibitor, verteporfin. Verteporfin is a suppressor of the YAP/TAZ-TEAD complex and activator of 14-3-3 that sequesters TAZ in the cytoplasm^[59]^. Treatment of NK cells with verteporfin enhanced NK cell cytotoxicity against K562 cells (**Figure 5D**) and upregulated the production of granzyme and perforin (**Figure 5E**). Additionally, verteporfin reduced KIR3DL1 surface expression (**Figure 5F**, red box), whereas PD-1 expression was unaffected. These observations suggest that one of the possible ways TAZ limits NK cell cytotoxicity is through the sustenance of inhibitory receptor, KIR3DL1.

**Figure 5.**
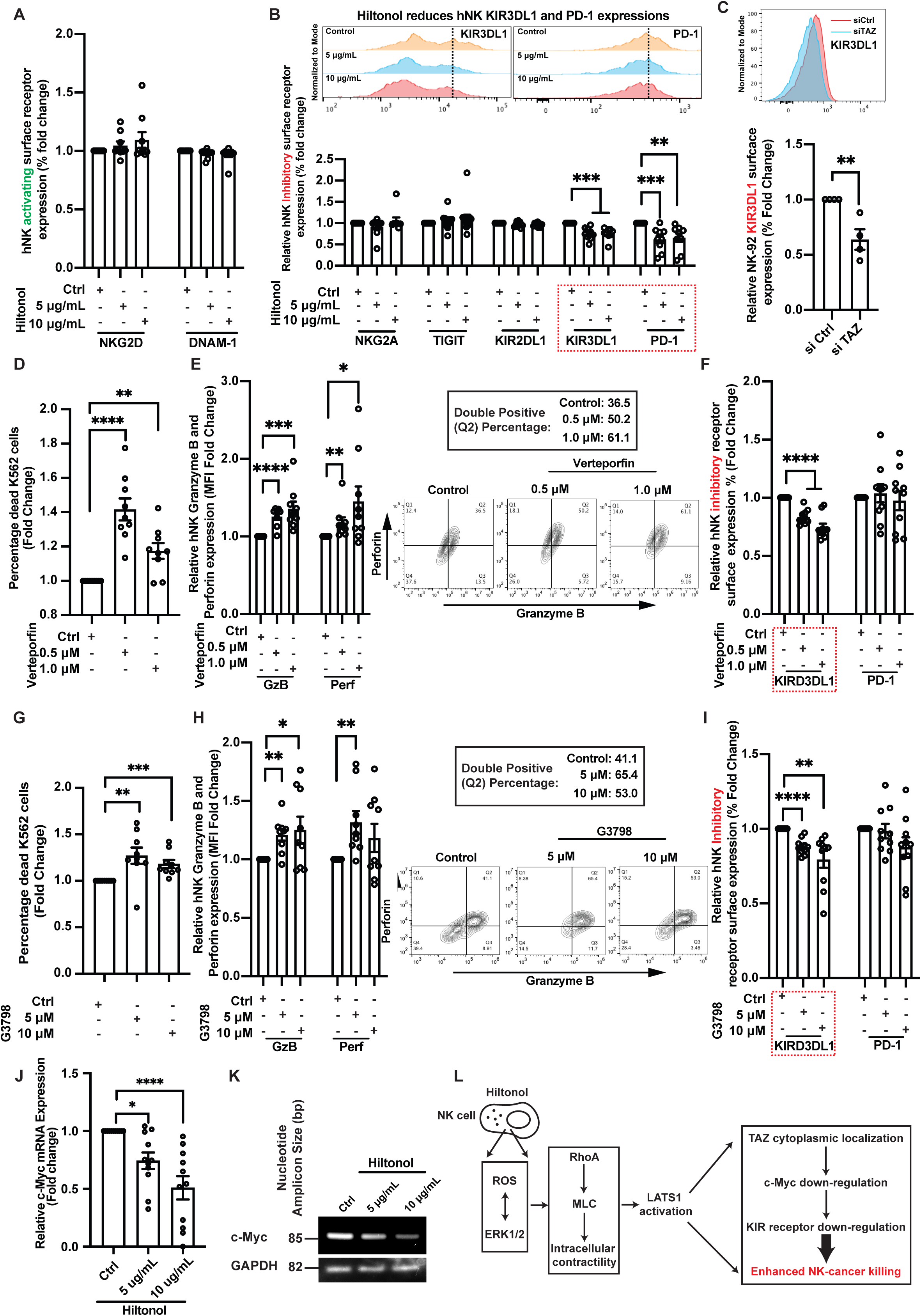
The NK cell inhibitory receptor, KIR3DL1, is regulated by TAZ-induced c-Myc activity. **(A) and (B)** Hiltonol treatment overnight (18 h) did not alter surface expressions of hNK activating receptors, NKG2D and DNAM-1 (A), whereas the inhibitory receptors KIR3DL1 and PD-1 were significantly reduced with hiltonol treatment (B), red box. The graph represents means ± SEM and n= 9 representative of three hNK donors. The representative flow cytometric histograms for KIR3DL1 and PD-1 for one donor is represented above the graph and the dotted vertical line facilitates the viewing of histogram shifts. **(C)** TAZ knockdown (48 h) in NK-92 cells significantly reduces the inhibitory receptor KIR3DL1 surface expression, whereas PD-1 surface expression is not affected. The graph represents means ± SEM and n= 4. **(D-F)** Overnight (18 h) treatment with YAP/TAZ inhibitor, verteporfin, increases hNK cytotoxicity against K562 lymphoblasts (D), with concomitant increase in expressions of granzyme B and perforin (E). On the other hand, only KIR3DL1 (red box) decreased with verteporfin treatment (F). The graph represents means ± SEM and n= 9 representative of three hNK donors. The flow cytometric layouts represent the increased granzyme B and perforin expression in hNK cell treated with verteporfin **(G-I)** The c-Myc inhibitor, G3798, treatment overnight (18 h) increases hNK cytotoxicity against K562 lymphoblast (G), with corresponding increase in the expressions of granzyme B and perforin (H). On the other hand, only KIR3DL1 (red box) decreases with G3798 treatment (I). The graph represents means ± SEM and n= 9 representative of three hNK donors. The flow cytometric layouts represent the increased granzyme B and perforin expression in hNK cell treated with G3798 **(J and K)** Hiltonol treatment reduces c-Myc mRNA expression. The graph represents means ± SEM and n= 11 representative of four hNK donors. A representative gel blot (out of n=4) was shown and the size markers represent the nucleotide amplicon size and not gene size. **(L)** Schematic showing the mechanistic enhancement of NK cell cancer killing through ROS-, ERK1/2-induced RhoA-MLC enhancement of intracellular contractility. Enhanced intracellular contractility activates LATS1 to inactivate TAZ through cytoplasmic localization that resulted in down-regulation of the inhibitory KIR receptor. For all graphs, *p < 0.05, **p < 0.01. ***p<0.001 and ****p < 0.0001 and One-way ANOVA was used to compare the different conditions.

Next, we verified if c-Myc, a downstream transcription factor under TAZ control, could have an influence on KIR3DL1. To this end, we tested whether c-Myc inhibition would enhance NK cytotoxicity. We found that inhibition of c-Myc with its inhibitor, G3798, increased NK cytotoxicity against K562 cells in a dose-dependent manner (**Figure 5G**). However, at a high dosage (20 μM), G3798 reduced the cytotoxicity of NK cells (data not shown). Furthermore, low doses of G3798 also resulted in an increase in granzyme B and perforin levels (**Figure 5H**). In addition, c-Myc inhibition reduced KIR3DL1 surface expression, without affecting PD-1 surface expression (**Figure 5I, red box**). This is supported by a previous report that c-Myc binding at the distal KIR promoter during NK-cell development specifically promotes KIR transcription^[60]^. Finally, treatment of NK cells with hiltonol was found to directly reduce mRNA transcript levels of c-Myc (**Figure 5J, K**), suggesting that hiltonol-induced TAZ cytoplasmic compartmentalisation resulted in and correlated with reduced c-Myc activity.

Overall, we have shown that TAZ in NK cells positively regulates c-Myc expression and function, which is modulated by hiltonol-induced intracellular contractility (**Figure 5L**). The expression of c-Myc sustains the expression of the inhibitory KIR3DL1 surface receptor. Interestingly, the KIR2DL1 inhibitory receptor was not affected (Figure 5B), possibly due to its very low expression on hNK cells. Hence, the regulation of KIR3DL1 surface expression by hiltonol could serves as a possible mechanism to regulate hiltonol-induced NK cell cytotoxicity.

### 2.6 Hiltonol treatment and c-Myc inhibition enhance the cytotoxicity of NK cells against breast and lung cancer cells

As a more stable form of dsRNA, Hiltonol has been widely used as a vaccine adjuvant^[19][61]^. Few studies have demonstrated direct cancer-killing capabilities of hiltonol on cancer cells^[42][43]^. However, whether hiltonol can participate in NK cell immunomodulation is unexplored. We have shown that hiltonol enhanced NK cytotoxicity, killing the model cell line, K562 lymphoblast (Figure 2I). Since we have previously demonstrated a graded NK cell response towards the less and more metastatic lung and breast cancer cells^[1]^, here we set out to determine if in coculture settings, hiltonol would enhance the cytotoxicity of NK cells towards the more resistant cancer types (lung and breast cancer) and subtypes (metastatic vs non-metastatic). Furthermore, metastatic lung and breast cancer patient and cells present high HLA ligand expression, and NK cells expresses high level of KIR3DL1 ^[62][63]^. This will provide a direct translational immunotherapeutic advancement.

We pre-treated hNK cells overnight (18 h) with hiltonol and then cocultured hNK cells with lung (H1299 and H1975) and breast (MCF7 and MDA-MB-231) cancer cells (**Figure 6A**). An overnight treatment with hiltonol was based on our previous observation that such a duration of treatment was sufficient to induce a robust early cytotoxic response in NK cells which correlated with transcription factor (Eomes) nuclear translocation. Furthermore, the less metastatic H1975 lung cancer and MCF7 breast cancer cell lines were easily killed by hNK cells^[1]^. We found that pre-treatment with hiltonol enhanced the cytotoxicity of hNK cells against both the breast and lung cancer cells (**Figure 6B i, ii**) by about 1.5 to 2.0 folds. Interestingly, this increase remained pronounced even when hNK cells were challenged with the metastatic cancer subtypes (MDA-MB-231 and H1299), which we have previously demonstrated to induce higher and prolonged contractility in NK cells. Consistently, the levels of perforin and granzyme B were increased (**Figure 6B iii, iv**). Nevertheless, the highly metastatic MDA-MB-231 could not induce an increase in hNK perforin (Figure 6B iv) despite hNK cells displaying higher killing capacity with hiltonol treatment (Figure 6B ii). Hence, other mechanisms may be involved to upregulate hNK-cancer cell killing.

**Figure 6.**
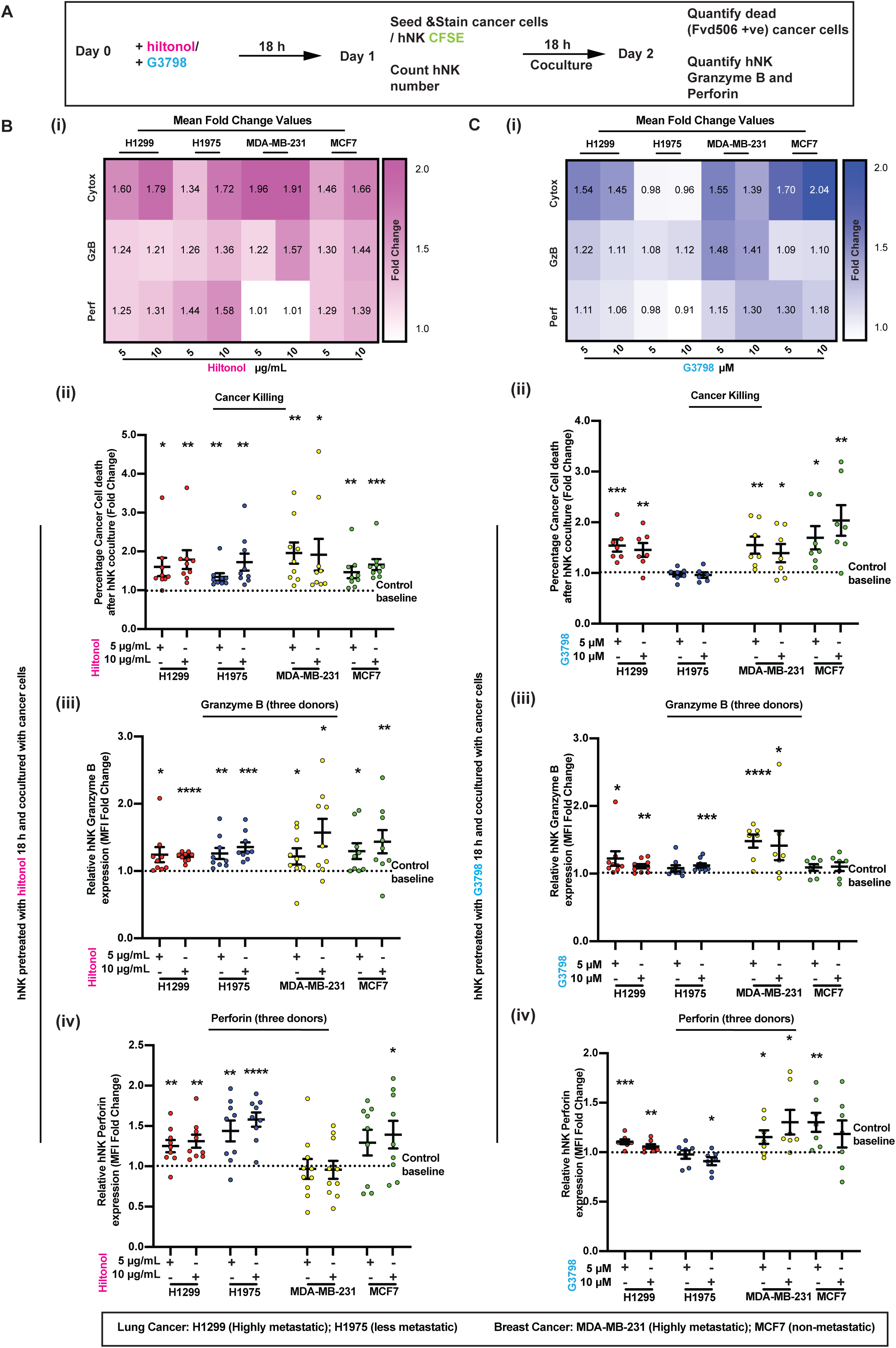
Hiltonol treatment and c-Myc inhibition enhance cytotoxicity of NK cells against metastatic and less metastatic breast and lung cancer cells. **(A)** Schematic flow showing the pre-treatment of hNK cells with hiltonol or G3798 overnight (18 h) before coculturing with cancer cells (seeded for at least 8 hours) to analyse cancer cell death or hNK granzyme B and perforin production. The cell type to be analysed was labelled with CFSE to allow easy differentiation during flow cytometric analysis and the FVD506 dye was used to stain dead cells. Refer to Figure S2 for gating strategy to identify CFSE staining dead cancer cells. **(B)** Hiltonol pre-treatment increases hNK cytotoxicity towards all lung (H1299 and H1975) and breast (MCF7 and MDA-MB-231) cancer cell subtypes (i) Heatmap showing the increase (fold change) in hNK cytotoxicity (Cytox), granzyme B (GzB) and perforin (Perf) against all cancer cell lines (Lung cancer: H1299 and H1975; Breast cancer: MDA-MB-231 and MCF7) with hiltonol pretreatment (5 and 10 μg/mL). The values in each of the heatmap cells are the mean values of data for ii-iv. (ii) Hiltonol treatment increases hNK cytotoxicity against all four cancer cell lines. The dotted horizontal line represents control baseline killing of hNK cells against the four cancer cell lines and is normalised to 1.0. Comparisons are made between each column to the control baseline. The graph represent means ± SEM and n ≥ 9, representative of three hNK donors. (iii and iv) Hiltonol pre-treatment increases hNK granzyme b (iii) and perforin (iv) production in a dose-dependent manner, by about 50%. Lung cancer cells (H1299 and H1975) induced higher hNK perforin production, while breast cancer cells (MCF7 and MDA-MB-231) induced higher hNK granzyme b levels. The dotted horizontal line represents control baseline killing of hNK cells against the four cancer cell lines and is normalised to 1.0. Comparisons are made between each column to the control baseline. The graphs represent means ± SEM and n ≥ 9, representative of three hNK donors. **(C)** The c-Myc inhibitor, G3798, enhanced or sustained hNK-induced cancer cell death in all cancer subtypes. (i) Heatmap showing the increase (fold change) in hNK cytotoxicity (Cytox), granzyme B (GzB) and perforin (Perf) against all for cancer cell lines (Lung cancer: H1299 and H1975; Breast cancer: MDA-MB-231 and MCF7) with G3798 pretreatment (5 and 10 μM). The values in each of the heatmap cells are the mean values of data for ii-iv. **(E)** Hiltonol was able to enhance more cytotoxicity against breast cancer cells (MCF7 and MDA-MB-231) and induced higher granzyme B and perforin levels in hNK cells (iii and iv). The dotted horizontal line represents control baseline killing of hNK cells against the four cancer cell lines and is normalised to 1.0. Comparisons are made between each column to the control baseline. The graphs represent means ± SEM and n≥ 6 representative of three hNK donors. For all graphs, *p < 0.05, **p < 0.01. ***p<0.001 and ****p < 0.0001 and One-way ANOVA was used to compare the different conditions.

Over the years, c-Myc inhibition has become the most appealing anticancer approach in cancer clinical trials. The expression of c-Myc is tightly controlled in normal cells, but becomes dysregulated and overexpressed in most human cancers, making it one of the most important human oncogenes^[64]^. However, *in vivo* observations often fall short of *in vitro* expectation^[64]^. Since c-Myc is one of the limiting factors in NK cytotoxicity (Figure 5), we cocultured hNK cells and cancer cells in the presence of c-Myc inhibitor (G3798), anticipating c-Myc inhibition to also increase NK cytotoxicity against the cancer subtypes.

Since c-Myc inhibition could induce inherent changes in hNK and cancer cell phenotype, we analysed the phenotype of hNK cells after G3798 treatment and showed that CD56 and CD16 surface expressions were not significantly affected (**Figure S4**). Next, we verified if c-Myc inhibition will result in direct cancer cell death. Interestingly, c-Myc inhibition did not result in cancer cell death unless it was applied at a high dose of 20 μM (**Figure S5A**) and the highly metastatic H1299 lung cancer cell line was most resistant to c-Myc inhibition. Conversely, c-Myc inhibition did not result in substantial cell death in NK cells, even at a high dose of 20 μM G3798 (**Figure S5B**), indicating the robustness of NKs. Since cancer cells were only susceptible to c-Myc inhibitor at a high dose (Figure S5A) and may present general toxicity for translational studies, we next cocultured hNK cells and cancer cells with G3798 at lower doses (5 and 10 μM) to investigate the specific impact of c-Myc inhibition of hNK on its anti-cancer activity. We found that c-Myc inhibition induced a dose-responsive increase in hNK-mediated cancer-killing capacity (**Figure 6C i, ii**). Strikingly, even the metastatic cancer subtypes (H1299 and MDA-MB-231) were susceptible to G3798-induced hNK killing. In addition, the increase in cancer killing capacity also correlated with a moderately increased granzyme and perforin levels (**Figure 6C iii, iv**). Therefore, our results agree with a previous study that c-Myc blockade with inhibitors suppresses cancer growth and this is further enhanced with combinatorial treatment with anti-PD-1 antibody^[65]^. Nevertheless, our observation that G3798 treatment induced only a modest increase in granzyme B and perforin production again implicates additional mechanism of hNK-induced cancer killing that remains to be explored.

Finally, to test whether hiltonol pre-treatment and c-Myc inhibition can elicit a synergistic effect on cancer-killing, we pre-treated hNK cells overnight with hiltonol before coculturing with highly metastatic cancer subtypes (MDA-MB-231 and H1299) in the presence of G3798. Interestingly, an apparent synergistic effect was observed when hNK cells were challenged with MDA-MB-231 cells **(Figure S5C).** This effect was less evident with the more resistant H1299 lung cancer cells. Since c-Myc is identified as a downstream target influenced by hiltonol treatment, our findings showed amplification (synergy) of the benefits of c-Myc inhibition with hiltonol-mediated antitumorigenic mechanisms^[17][43][66]–[68]^.

In conclusion, we have shown that the innate immune NK cells express TAZ but not YAP. The activation of intracellular contractility by the immune adjuvant, hiltonol, through ERK1/2 activation and ROS production led to TAZ cytoplasmic compartmentalisation and enhanced NK cytotoxicity. This was achieved by release of LATS1 from F-actin filaments which correlates with increased LATS1 activation and TAZ inactivation. In the absence of hiltonol, homeostatic control of contractile forces maintains c-Myc activity, resulting in the surface expression of the inhibitory receptor KIR3DL1. Our findings provide an avenue for therapeutic translational advances as our results indicate that sensitising hNK cells to hiltonol could subdue both metastatic and non-metastatic cancer types (breast and lung) and the corresponding subtypes. **Figure 7** provides a model to summarize our findings.

**Figure 7.**
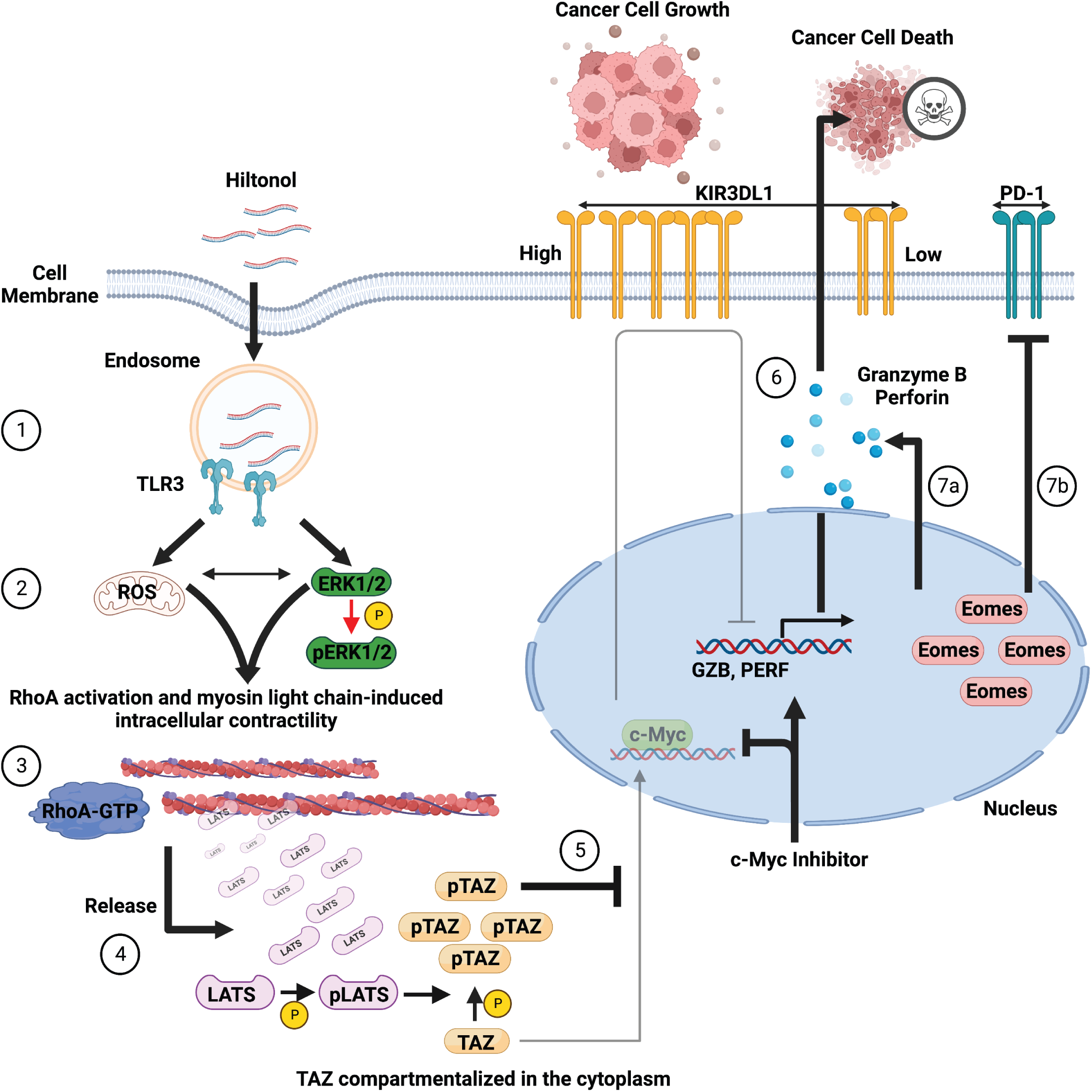
Schematic showing hiltonol treatment enhanced NK anti-cancer capacity. **(1)** Hiltonol increases NK cell TLR3 expression that correlated with enhanced ROS (mitochondrial and cytoplasmic) and ERK activation. **(2**) ROS and ERK activation form a positive feedback loop. **(3)** ERK activation resulted in RhoA activation and enhanced myosin light chain-based intracellular contractility, resulting in **(4)** the release of LATS1 from actin. This correlated with LATS1 activation through phosphorylation, which inactivates TAZ through phosphorylation. **(5 and 6)** Phosphorylated TAZ is unable to enter the nucleus and induce c-Myc activity associated with increased KIR3DL1 expression. This was supported by c-Myc inhibition (with G3798), which resulted in increased expression of GZB (granzyme b) and PERF (perforin). **(7a and b)** Enhanced contractility also resulted in Eomes nuclear localization and this was associated with reduced PD-1 expression and increased granzyme b and perforin production. Altogether, NK cells treated with hiltonol and c-Myc inhibitor display enhanced cytotoxicity against breast and lung cancer cells.

## 3. Discussion

The roles of cell ‘force’ in regulating pathophysiological processes are gaining attention and warrant urgent attention. Various NK cell-related immune diseases result from aberrant regulation of proteins that control cytoskeletal force^[27]^. Here we provide empirical evidence supporting NK cell intracellular contractility modulated with an immune adjuvant, hiltonol. The direct consequence of this is the cytoplasmic sequestration with phosphorylation and inactivation of TAZ protein, and subsequent down-expression of the surface inhibitory receptor, KIR3DL1. Interestingly, we also observed an indirect down-regulation of an important inhibitory receptor, PD-1, that was associated with an increase in Eomes nuclear localization. Surmise to say, pre-treatment of NK cells with hiltonol can upregulate the cytotoxicity of NK cells to target and kill lymphoblast and breast and lung cancer cells.

The mechanistic role of ‘force’ in triggering YAP/TAZ localization in cell systems is gaining attention since various studies have shown that force does not seem to act alone in this process. It was previously demonstrated that force applied on the nucleus was sufficient to trigger YAP nuclear localization^[33]^. We have previously shown that the transcription factor, Eomes, entered the NK nucleus as a consequence of increased intracellular contractility^[1]^. Although YAP/TAZ can be regulated canonically by a series of kinases (MST1/2 and LATS1/2)^[29]^, how these kinases are regulated by cell force is unclear. YAP has been shown to suppress T-cell activation and differentiation^[34][35][69]^, but we have shown that NK cells express TAZ, but not YAP. With the enhancement of RhoA-MLC2 induced intracellular contractile force, TAZ was found to be sequestered in the cytoplasm instead of entering the nucleus. Our findings agree with other studies whereby force alone may not lead to YAP nuclear localization. In a study with gastric cancer cells, the presence of cadherin 1 was purportedly required for YAP nuclear localization as a result of gain-of-function RhoA activation^[70]^. In yet another cardiomyoblast differentiation model, intracellular contractile force was shown to sequester YAP in the cytoplasm to switch the cellular process from growth to differentiation^[71]^. Here, we show that ‘force’, promotes the cytoplasmic sequestration of TAZ in NK cells. Importantly, our results provided mechanistic explanations on how force activates the inactivating kinase, LATS1, to inactivate TAZ. Since LATS1 is known to bind to actin^[55]^, the conceptual understanding is that we view actin as a relaxed string with LATS1 bound to it. We perceive that activation of force through RhoA-induced MLC2 activation creates ‘tension’ on the actin ‘string’ that results in the ‘release’ of LATS1 from actin and this event correlated with LATS1 activation through phosphorylation (see Figure 4C). At this juncture, we do not yet have an explanation on how the release of LATS1 from actin results in its activation. However, it is conceivable that the binding of LATS1 to actin directly resulted in a ‘microenvironment’ that created intense competition for the abundance of ATP available for activation of actin and LATS1. Actin is well known to bind ATP or ADP monomers tightly^[72]^. Two studies with small inhibitors of LATS1 also demonstrated that competition for ATP inhibits LATS1 activation^[73][74]^. Hence, it is likely that the ‘microenvironment’ consisting of actin and LATS1 favours actin activation. The dissociation of LATS1 from actin may expose LATS1 ATP-binding sites and facilitate its activation through phosphorylation (Figure 4D). On the other hand, the activation of LATS1 following its dissociation from actin may depend on other scaffold proteins that also bind to actin (e.g. angiomotins). It is known that the scaffold protein angiomotin and LATS1 are able to activate each other^[75][76]^. Importantly, the mutant defective in angiomotin which was unable to bind F-actin, was shown to inactivate YAP in various cell models and the deletion of LATS1 had an additive effect^[77]^. Hence, it is possible that other components of Hippo-TAZ signaling pathway may be regulated by enhancement of intracellular contractility through dissociation from actin. Nevertheless, whether and how LATS1 and actin compete for ATP, and whether the LATS1 activation through enhanced contractility requires other kinases that bind to actin, warrants further investigation. Furthermore, whether the dynamics of LATS1 recruitment to actin is affected by contractility could also be studied in detail in future by using various high resolution and quantitative microscopy techniques (e.g. Fluorescence Recovery After Photobleaching).

TLR3 is a well-studied PRR that activates anti-viral responses in NK cells^[17]^. However, what is unclear is how TLR3 may activate contractile forces in a context dependent manner. For instance, TLR3 activation by Poly-I:C activated contractility in aortic smooth muscle cells^[22]^. Partly inspired by this, we investigated the possibility that NK cells, which lack the force-transducing focal adhesion complex, can still activate contractility through TLR3 activation. Interestingly, we showed that hiltonol was able to increase the contractile force of NK cells, and this was through the activation of RhoA and MLC2. In various cellular and cancer models, TLR3 was also shown to activate small GTPases like RhoA, Cdc42 and Rac1^[78]–[80]^. Importantly, what are the functional consequences of enhanced contractility? Although we have shown that TAZ cytosolic localization and Eomes nuclear localization govern the increase in NK cytotoxicity, we do not discount the possibility that various pathways can be activated by contractile forces or through RhoA that is independent of force. For instance, it has been demonstrated that RhoA is required for NF-κB activation^[78]^ and NF-κB itself is regulated by TLR3 activation and has been shown to promote NK anticancer activities. Interestingly, NK cell activation also depends on other immune subtypes *in vivo*. For instance, Poly-I:C was shown to activate dendritic cells that subsequently activated NK cells^[66]^; and the activation of NF-κB was shown to facilitate NK activation by dendritic cells^[81]^. Importantly, apart from TLR3, other intracellular dsRNA sensors, MDA-5 and RIG-1, are also reported to respond to Poly-I:C in mouse dendritic cell activation^[67]^ and this may cross-activate NK cells. Hence, hiltonol administration *in vivo* may activate contractility via multiple PRRs in multiple immune cell types and together, upregulate anti-cancer activities. Future studies may be required to distinguish distinct and complementary roles of the various PRRs in response to hiltonol pre-treatment in humans *in vivo*. After all, mouse NK cells are distinctive from human NK cells. For instance, we showed previously that hNK cells express similar levels of the transcription factor Eomes after activation, whereas murine NK cells showed differential Eomes expression^[1]^. Nevertheless, NK cells are the sentinel defenders against pathogens and cancers.

In this study, we utilised various small molecule inhibitors to pharmacologically inhibit TAZ activity (verteporfin) and c-Myc (G3798) activity. The inhibition of c-Myc, a downstream target of the transcriptional co-activator TAZ, likewise enhanced the cytotoxicity of NK cells against the cancer cell lines studied. However, inhibition of c-Myc seemed to result in lesser cancer cell death (∼ 1.5-fold) compared to hiltonol treatment (∼ 2.0-fold). These observations again suggest that hiltonol inactivation of TAZ through RhoA-MLC2 and LATS1 activation may not be the only signaling axis. As suggested above, other pathways may be simultaneously activated by TLR3 activation. On the other hand, our results vividly illustrate how the subcellular localization of two endogenous proteins, TAZ and Eomes, affects NK cytotoxicity, which could be potentiated by an exogenous chemical inhibitor of c-Myc, G3798. It would be pertinent in the future to test whether a cocktail of various inhibitors, activators and neutralizing antibodies would further enhance NK cell cytotoxicity. Some of the options may include c-Myc inhibitor combined with verteporfin or hiltonol, hiltonol combined with IL15 and other endogenous proteins such as TGFβ that activates NK contractility^[1]^. However, a note of caution is a possible increase in ROS generated by NK cells, to levels which may be self-cytotoxic. We showed that a high dosage of hiltonol (25 μg/mL) increased ROS production which reduced the enhanced NK cytotoxicity. A small amount of ROS was proficient in improving NK metabolism and function^[47][51][52]^. However, high ROS levels is cytotoxic. Our findings may perhaps explain why high doses of hiltonol/ Poly-I:C were unable to sustain an increase in NK cell cytotoxicity *in vitro* and *in vivo*^[41]–[43][82]^.

Altogether, we have demonstrated a mechanistic understanding on how cell ‘force’ (intracellular contractility) inactivates TAZ to promote NK cytotoxicity towards cancer cells. Our findings may provide an innovative avenue for *ex vivo* rejuvenation/activation of autologous NK cells for anti-cancer immunotherapies.

## 4. Experimental Section

Except for Imaging flow cytometry, proximity ligation assay and DHE and MitoSOX assays, all other methodologies are adapted from our previous publication^[1]^ with slight modifications. The resource table for antibodies and primers, and flow cytometry gating and controls used in this paper can be found in the supplementary material (**Supplementary Table S1, S2**).

### Cell culture and drug treatments

Isolated primary human NK (hNK) cells were expanded for one week before experiments following the manufacturer’s recommendation. hNK was cultured in NK MACS medium (Miltenyi Biotec) supplemented with 1% NK MACS (Miltenyi Biotec), 5% human AB serum (Sigma-Aldrich) and 25 ng/mL IL-2 (Miltenyi Biotec). NCI-H1299, NCI-H1975, MCF7, MDA-MB-231 and K562 were obtained from the American Type Culture Collection. NK-92 cell line was maintained in RPMI 1640 supplemented with 12.5% FBS, 12.5% horse serum (Gibco), 1% penicillin/streptomycin and supplemented with 10 ng/mL IL-2. For non-small cell lung cancer cells, NSCLCs (NCI-H1299 and NCI-H1975) and breast cancer cell lines (MCF7 and MDA-MB-231), TrypLE Select Enzyme (Gibco) was used for passaging cells. Only cells within passages 3 to 20 were used for experiments. All cell lines were supplemented with Plasmocin Prophylatic (Invivogen™ Cat# ant-mpp) for the prevention of mycoplasma contamination and routinely tested and verified to be free from mycoplasma using MycoStrip (Invivogen™ Cat # rep-mys) (**Supplementary Figure S8**). All cells were grown at 37°C with 5% CO2.

Poly:ICLC (Hiltonol) is a clinical grade formulation of poly-I:C stabilized with poly-L-lysine and carboxymethylcellulose manufactured by Oncovir. For drug treatments, cultured cells were treated with hiltonol (Oncovir Inc.), c-Myc inhibitor G3798 (Sigma-Aldrich, Cat# 10074-G5), verteporfin (Sigma-Aldrich, Cat#SML0534) and blebbistatin (Sigma-Aldrich, Cat# 203390) at indicated concentrations represented on the figures. 1X PBS and DMSO were used as vehicle controls for hiltonol, and G3798, verteporfin and blebbistatin respectively. The Rho activator CN03 was purchased from Cytoskeleton Inc. (Cat# CN-03) and diluted in sterile water.

### Isolation of PBMC and primary NK cells

The protocols for isolation of PBMC from healthy donors were approved by the Health Sciences Authority Singapore (HSA) and the National University of Singapore Institutional Review Board (201706-06 and H-17-028E). Apheresis cones from healthy donors were obtained from the HSA. Peripheral blood mononuclear cells (PBMCs) were isolated from the blood samples using Ficoll-Paque (GE Healthcare, Cat# 17144003) and gradient centrifugation. The middle layer containing PBMCs were then enriched for primary human NK cells (hNK) by negative selection using the EasySep Human NK Cell Enrichment Kit (StemCell Technologies, Cat# 19055), following the manufacturer’s protocol. Isolated hNK cells are CD56+ and CD3- (**Supplementary Figure S7A)**

### Coculture of NK cell with lung or breast cancer cells

The effector (NK-92 or hNK) to target (cancer cells) were cocultured in a ratio of 2.5:1 as previously determined^[1][83]^. Briefly, hNK and NK-92 cells were pre-stained with CFSE (Sigma) according to the manufacturer’s protocol, which was able to perpetuate stained cells for as long as 6 days as determined previously^[1]^, except for those used for cytotoxicity assay against model target cell line, K562. Lung and breast cancer cells were pre-seeded for at least eight hours for them to attach to cell culture plate. Coculture was carried out in NK MACS media supplemented with 25 ng/mL IL2 for hNK cells, or 10% RPMI supplemented with 10 ng/mL IL-2 for NK-92 cells.

### NK cell cytotoxicity assay against K562 cells

The effector (NK-92 or hNK) to target (K562) was 5:1 and 1:1, respectively, as previously determined by the group^[1][83]^. Target cells were pre-stained with CFSE prior to coculture with NK cells for cytotoxicity assay. After a 4-hour incubation period maintained at 37°C with 5% CO2, dead cells were stained with fixable viability dye eFluor506 (eBioscience, Cat# 65-0866-14) on ice for 30 minutes and immediately fixed with 2% PFA for analysis using flow cytometry. The gating strategy to identify CFSE-labelled dead K562 cells are in **Figure S2**.

### Transfection and transduction (siRNA and lenti-virus)

NK cells were transfected with scrambled or ON-TARGET plus human WWTR1 SMARTpool siRNA from Dharmacon Inc, (CCGCAGGGCUCAUGAGUAU, GGACAAACACCCAUGAACA, AGGAACAAACGUUGACUUA, CCAAAUCUCGUGAUGAAUC) using the Neon Transfection System (Invitrogen) and according to the manufacturer recommended parameters. Following transfection, cells were allowed to recover in antibiotic-free media for 48 hours.

The Lenti-ORF clone of Human WW domain containing transcription regulator 1 (WWTR1/ TAZ), transcript variant 1, mGFP-tagged, in vector pLenti-C-mGFP-P2A-Puro was purchased from manufacturer, OriGene (Cat# RC231269L4). The manufacturer’s protocol was used for viral production by mixing pLenti-C-TAZ-mGFP-P2A-Puro with packaging plasmids and transfection reagent from the Lenti-vpak packaging kit (OriGene, Cat#TR30037). The mixture was transfected into HEK293T cells to generate pseudoviral particles. NK-92 cells stably expressing TAZ-mGFP were established by transduction with pseudoviral particles with 8μg/mL polybrene (Sigma-Aldrich, Cat# TR-1003-G). Stable cell lines were established through selection with puromycin (Sigma-Aldrich, Cat# P4512) and fluorescence-activated cell sorting.

### Western blot analysis and active RhoA binding assay

The Western blot procedures were performed, as previously described^[1]^. Briefly, cells were washed once with ice cold 1X PBS and lysed with ice cold RIPA supplemented with protease inhibitor, sodium orthovanadate, sodium fluoride and β-Glycerophosphate. BCA assay was performed to prepare samples of equal protein concentration. 4X loading dye containing 2-Mercaptoethanol was added to the sample before boiling the sample at 85 °C for 10 minutes and ran on a denaturing SDS-PAGE gel. The results were visualised by ChemiDoc Touch (Bio-Rad) and analysed using Image Lab V 5.2.1. The antibodies used are listed in **Supplementary Table S2**.

The Rhotekin-RBD Beads (Binds Active Rho Proteins) from Cytoskeleton Inc. (Cat#RT02) was used to pull down active RhoA-GTP in cell lysates. The procedures for pull down follows the manufacturer’s protocol.

### RT-qPCR analysis

RNA extraction was carried out using Pure-NA™ Fast Total RNA extraction kit following the manufacturer’s protocol. The cDNA across samples was generated from equal amounts of RNA using SuperScript IV VILO Master Mix following the manufacturer’s protocol. Diluted cDNA (20 ng/ul) was used to quantify the delta Cq values of the samples using SsoFast™ EvaGreen® Supermix from Biorad on the Bio-Rad CFX 96 Real-Time PCR Detection System.

### Immunofluorescence and microscopy

For immunofluorescence staining, cells were centrifuged at 500 xg for 5 minutes and fixed with 4% PFA for 15 min at 37°C. Free aldehydes were quenched with freshly prepared sodium borohydride (0.01%; Sigma-Aldrich, Cat# 452882) dissolved in 1x PBS for 5 min. Samples were washed thrice with 1x PBS at 5-minute intervals and 3% bovine serum albumin and 0.2% Triton X-100 in 1× PBS (blocking buffer) was used to permeabilize and block the samples. The appropriate primary antibodies were dissolved in blocking buffer following manufacturer’s recommendation, and incubated for 45 minutes at room temperature. This was followed by washing thrice with 1× PBS and incubating with Goat anti-Rabbit IgG (H+L) Highly Cross-Adsorbed Secondary Antibody, Alexa Fluor– 647 conjugated secondary antibodies (ThermoFisher Scientific, Cat# A32733), and DAPI (ThermoFisher Scientific, Cat# D1306) diluted in 3% BSA in 1X PBS. Finally, the cells were washed thrice with 1× PBS and placed in an iBidi glass-bottom dish (iBidi, Cat# 81218-200) pre-coated with 0.01% poly-L-lysine (Invitrogen, Cat# P8920). The glass bottom dishes were centrifuged at 500 xg for 5 minutes using a swing-out bucket rotor. For visualisation of actin, Alexa Fluor 488 Phalloidin (ThermoFisher Scientific, Cat# A12379) stain was used together with the secondary antibodies at 1:500 dilution.

Imaging was performed on a Yokogawa CSU-W1 (Nikon TiE system) spinning disk microscope using a 1.40 numerical aperture (NA) oil immersion 100x objective. For comparison between samples, the binning, laser powers and exposure time were kept constant, and all samples were stained with a master mix of primary and secondary antibodies.

### Laser ablation of NK cell actin filaments

The recoil velocity (measured by laser ablation) due to tensional release of actin network mesh within the cells is measured by monitoring distinctive features before, during and after UV laser ablation. Spatiotemporal information of the distinctive features after UV ablation was obtained and quantified using MTrackJ plugin in Fiji imageJ. Fitting of calculated distance between points of distinctive features was carried out using a linear function and a single exponential function, or a linear function and a double exponential function. The single exponential function was used for datasets with slow recoil velocity to obtain slow elastic response of the actin network mesh upon ablation. To obtain datasets with fast recoil velocity, double exponential function was used for fast elastic response of the actin network mesh upon ablation. Recoil velocity was calculated as the derivative of the abovementioned functions using methods mentioned in earlier literatures^[1][46][84]^.

The linear function is as shown,

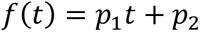

Where ***t*** is approximate time when UV laser shutter is opened, ***p***_***1***_ is deformation speed for linear model, and ***p***_***2***_ is the initial length between points of distinctive features at t=0 s for linear model.

The single exponential model is as shown,

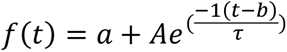

Where ***t*** is approximate time when UV laser shutter is opened, ***a*** is the initial length between points of distinctive features at ***t*** = 0 *s* for exponential model, ***τ*** is ratio of Young’s modulus to viscosity, and ***A***, and ***b*** are arbitrary constants.

The double exponential model is as shown,

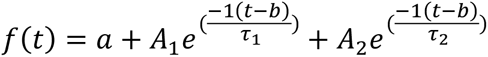

Where ***t*** is approximate time when UV laser shutter is opened, ***a*** is the initial length between points of distinctive features at ***t*** = 0 *s* for exponential model, ***τ***_***1***_ and ***τ***_***2***_ are ratios of Young’s modulus to viscosity, and ***A***_***1***_, ***A***_***2***_, and ***b*** are arbitrary constants.

### Flow cytometry and Imaging flow cytometry

The antibodies used for flow cytometry analysis are represented in **Table S2** and their respective FMO are represented in **Figure S7B**. For cell surface staining of receptors, fluorophore-conjugated antibodies were diluted in 1x PBS supplemented with 2% FBS at 1:100 dilution. NK cells were stained with the respective antibodies on ice for 10 minutes, followed by staining with fixable viability dye eFluor506 (eBioscience) on ice for 30 minutes. The cells were immediately sent for flow cytometry analysis to distinguish between live and dead cells and the expression of surface proteins.

To detect intracellular proteins, cells were first stained with fixable viability dye eFluor506 (eBioscience) on ice for 30 minutes and subsequently fixed with Intracellular Fixation and Permeabilization Buffer Set (eBioscience) for 40 minutes. Fixed cells were then stained for 30 minutes with the respective conjugated antibodies in permeabilization buffer at 1:100 dilution. Flow cytometry data were acquired using the CytoFLEX LX machine from Beckman Coulter, and all flow cytometry data were analysed using FlowJo V10.4.

For imaging flow cytometry, samples were prepared according to the immunofluorescence staining methodology, to stain for DAPI and TAZ. Data were obtained using Amnis® ImageStreamX Mark II Imaging Flow Cytometer. Fluid pressure was set to ‘low’ for high sensitivity. The cells were gated sequentially for high HMG (better focus), single cells, DAPI-TAZ double-positive cells, and nuclear localization (high similarity dilate). The template was saved and used for analysis of different sample conditions. A single colour control (DAPI or TAZ only) was used to automatically populate the compensation matrix. All data were analysed with the IDEAS 6.0 software using the wizard for nuclear translocation. DAPI (Ch7) was used as nuclear probe and TAZ (Ch11) was used as the translocating probe. The gating strategy is represented in **Figure S6**.

### Proximity ligation assay

Proximity ligation assay experiments were carried out using Duolink^®^ flowPLA Detection Kit - FarRed (Sigma Aldrich, # DUO94004) following the manufacturer’s protocol. A sample with no primary antibodies but with the PLA probes was used as the technical negative control as recommended by the manufacturer. NK cells were fixed with warm 37 °C PFA for 15 min at room temperature (RT) and subsequently permeabilized with 0.2 % Triton X-100. Afterwards the samples were blocked with Duolink^®^ Blocking Solution and incubated with primary antibodies (mouse actin and rabbit LATS1 antibodies; Table S2) for 1 hour. The subsequent steps of the proximity ligation assay (PLA) followed the manufacturer’s instructions. Data analysis of flow-cytometry PLA was performed using CytoFLEX LX machine from Beckman Coulter, and all flow cytometry data were analysed using FlowJo V10.4.

### DHE and MitoSOX assay

The Dihydroethidium (DHE) assay kit for the measurement of intracellular superoxide and hydrogen peroxide was purchased from Abcam (Cat#ab236206). The MitoSOX^TM^ for the measurement of mitochondria superoxide was purchased from ThermoFisher Scientific (Cat#M36008). The manufacturer’s protocol was followed throughout. Antimycin A and N-Acetyl Cysteine Assay from the DHE assay kit were used as positive and negative controls, respectively. All cells were analysed live and no fixation was involved in any step. Data acquisition of DHE and MitoSOX^TM^ signals were performed using CytoFLEX LX machine measured in PE channel, and all flow cytometry data were analysed using FlowJo V10.4.

### Quantification of nuclear-cytoplasmic ratio

Microscopy images obtained were analysed using Image J V2.0. The nuclear/ cytosolic ratio was analysed as previously described^[1][33]^, with slight modifications to accommodate the smaller cytoplasmic areas in NK cells. A Z-stacked image (step size 0.2 μm) encompassing the whole cell was obtained and the stacks demarcated by nuclear DAPI staining were Z-stacked for quantification of nuclear/cytoplasmic ratio. The DIC image, DAPI, and antibody channels were then merged to demarcate, respectively, the cell cytoplasm (CFSE and DIC areas without DAPI stain) and the nucleus (DAPI) and TAZ. To avoid measurement of an ROI (Region on Interest) in the nucleus that exceeded the cytoplasmic ROI, the biggest possible ROI drawn in the cytosol next to the DAPI-stained nucleus was first measured. The same ROI was then shifted inside the nucleus to measure the mean intensity of TAZ within the nucleus. The nuclear-cytoplasmic ratio was derived by dividing the nuclear intensity by cytosol intensity.

### Analysis of NK cell nuclear size and perimeter

To analyse the geometric properties of the nuclei of NK cells, DAPI-stained nuclei images were first segmented from the stack images before summing the z-projected slices to convert the x-y-z images into x-y images. To remove and blur out the speckled features within the nuclei, images were filtered with Gaussian Blur. Next, nuclei objects were separated from the background through adjusting the threshold to obtain the images in black and white (0-255). To separate nuclei touching each other, Watershed function built-in in ImageJ was utilized to find the centre of each nuclei object, distance map from nuclei object center points to edges of nuclei objects calculated, then a topological map was formed with lines created to separated nuclei touching one another. Once images were processed, Analyze Particles built-in function in ImageJ was used to locate the nuclei objects, count them, and obtain statistical parameters (e.g. perimeter of nucleus, area of nucleus) from the region of interest (ROI) demarcating each nucleus. Incomplete nuclei on the edges of the image were excluded.

### Statistical analysis

All graphs are represented as mean *±* SEM. Statistical analysis was carried out using Prism 8.2.1 (GraphPad Software). For statistical significance, p-value < 0.05 is considered as significant (*p < 0.05, **p < 0.01, ***p < 0.001, ****p < 0.000). Two-tailed Student’s T-tests were used when two cases were compared, and analysis of variance (ANOVA) test was used when more cases were analysed. The non-parametric tests were applied for data that did not meet normality criteria. For all experiments, donor hNK cells isolated from at least three different PBMCs were used. At least three replicates were performed with all other cell lines used in this paper.

### Data and materials availability

All data needed to evaluate the conclusions of the paper are present in the paper and/or the Supplementary Materials. Additional data and codes related to this paper may be requested from the authors upon reasonable request.

## Acknowledgements

This project is supported by the National Research Foundation, Singapore, under its funding to NUS for RIE-Related Roles for the SGUnited Jobs Initiative (NRF-MP-2020-0004). We thank the core facilities of the National University of Singapore, Department of Biological Sciences (NUS-DBS), Confocal Microscopy Unit and Flow Cytometry Laboratory (NUS-CMA), Department of Microbiology (NUS-Microbiology) and Mechanobiology Institute of Singapore, and Centre for BioImaging Sciences (NUS-MBI) for technical support. We thank Dongxue Hu for advice on DHE and MitoSOX assays. The illustrations were created with biorender.

## Author Contributions

D.W.C.P, Z.X, N.S, J.Y.Y, I.Y. and T.T conducted the experiments; A.M.S provided hiltonol and background information. Y.C.L, B.C.L and J.L.D provided advice and samples or reagents; D.W.C.P and J.L.D designed the experiments and wrote the manuscript. All authors commented on the manuscript.

