## Supplementary Figure Legends and Tables for "Natural Killer cell contractility and cytotoxicity is driven by TLR3 agonist-mediated TAZ cytoplasmic sequestration"

**Supplementary Information**

**Supplementary Figures**

**Supplementary Figure S1. NK cells express TAZ but not YAP. (A)** NK cells express moderate levels of TAZ (WWTR1) (highlighted in red) compared to other immune cell lineages. The graph represents the transcripts per million protein coding genes (pTPM) for each of the single cell clusters. Data are obtained from RNA-sequencing of 29 immune cell types from PBMCs of healthy donors[36]. **(B)** With the exception of basophils and progenitor cells, all other immune cell types do not express YAP mRNA. The graph represents the pTPM for each of the single cell clusters. Data are obtained from RNA-sequencing of 29 immune cell types from PBMCs of healthy donors[36].

**Supplementary Figure S2. Gating strategy for FVD506 staining of dead cells.** For the staining of dead cells for cytotoxicity assay or general protein expression quantification, the target cells are pre-stained with CFSE dye to allow flow cytometric differentiation. Subsequently, the cells are gated sequentially for differentiation from debris (FSC-H: SSC-H), single cells (FSC-H: FSC-A), CFSE positive cells (CFSE: FSC-A) and FVD506 positive (dead)/ negative (live) cells. The example shows the gating of dead CFSE-stained K562 cells that have been killed by hNK cells.

**Supplementary Figure S3. Hiltonol increases TLR3 expression, induces Eomes nuclear localization in NK cells and promotes NK: K562 binding. (A)** Hiltonol induces a dose dependent increase in hNK TLR3 expression. The graph represents means ± SEM, and n=9 representative of three hNK donors. **(B)** Hiltonol treatment does not cause spontaneous death in hNK cells. The graph represents means ± SEM, and n=9 representative of three hNK donors. **(C)** Hiltonol treatment overnight reduces hNK cell maximum projected nuclear area (C) and perimeter (D). The graphs represent mean ± SEM and, n= 148, 104, 131 for control (ctrl), 5 μg/ml and 10 μg/ml hiltonol, respectively, for both (C) and (D). The hNK cells analyzed are representative of three individual healthy donors. **(E)** Hiltonol treatment induces nuclear localization of Eomes in hNK cells. The dotted lines demarcate the cell periphery. Scale bar = 10 μm. The graph on the right represents means ± SEM, and n=42, 49, 40, for control (ctrl), 5 μg/ml and 10 μg/ml hiltonol, respectively. **(F and G)** Cell:cell interaction /binding assay shows that hiltonol treatment increases the binding percentage of CFSE labelled NK-92 (F) and hNK (G) cells to CellTrace™ Far Red labelled K562 cells. n=4 for NK-92 cells and n=9 for hNK cells, representative of three hNK donors. For all graphs, *$p < 0.05$, **$p < 0.01$, ***$p < 0.001$ and ****$p < 0.0001$ and One-way ANOVA was used to compare the different conditions.

**Supplementary Figure S4. Hiltonol and G3798 treatment do not alter hNK surface CD56 and CD16 expressions.** Representative flow cytometric plots of CD56 (x-axis) and CD16 (y-axis) of hNK cells from one donor treated with control (ctrl), 5 μg/ml and 10 μg/ml hiltonol (top) or control (ctrl), 5 μM and 10 μM G3798 (bottom).

**Supplementary Figure S5. High dose of c-Myc inhibitor (G3798) induces cell death on less metastatic cancer cells. (A)** G3798 treatment induced small amount of cell death in all cancer cell subtypes. However, the breast cancer cell lines (MCF7 and MDA-MB-231) were most susceptible to c-Myc induced cell death and peaked at 20 μM. The graph represents means ± SEM, and n=4 for all cancer cell lines. (B) G3798 treatment did not induce NK cell death even at a relatively high dose (20 μM). The graph represents means ± SEM, n=4 for NK-92 cells and n=9 for hNK cells, representative of three hNK donors. **(C)** Combinatorial treatment of hiltonol and G3798 induced greater breast cancer (MDA-MB-231) cell death compared to hiltonol or G3798 alone treatments. Lung cancer (H1299) showed a similar but less significant profile). The graph represents means ± SEM, n=6 for hNK cells, representative of three hNK donors. For all graphs, *p < 0.05, **p < 0.01. ***p<0.001 and ****p < 0.000, and One-way ANOVA was used to compare the different conditions. The two-tailed students *t-test* was used to compare between two groups.

**Supplementary Figure S6. Gating strategies for flow cytometry experiments used in the study. (A)** Gating strategy to obtain binding percentage of CFSE labelled NK cells to CellTrace™ Far-Red-APC K562 cells. Intact cells are first differentiated from the debris, and gated. Next, a quadrant (Q) is drawn and Q2 represents CFSE and Far-Red-APC double positive cells. The gating strategy is adopted from the Zheng et al. [85]. **(B)** Imaging flow cytometry gating was performed following the manufacture's protocol for nuclear translocation analysis. Cells in focus are first gated, followed by single cells and TAZ, DAPI double positive cells. R4 represents the relative TAZ and DAPI nuclear overlap values. The template is used for comparisons across different sample conditions to ensure same analysis is conserved. The similarity dilate score is tabulated from R4.

**Supplementary Figure S7. Flow cytometry antibody controls used in this study. (A)** hNK cells isolated from PBMCs using NK cell enrichment kit are CD56 positive (+) and CD3 (-) as per manufacturer's protocol. Please refer to materials and methods for detailed description on isolation. **(B)** The Fluorescence Minus One (FMO) or all conjugated antibodies for flow cytometry are represented in the figure. The blue line represents the fluorophore conjugated antibody and red line represents the FMO.

**Supplementary Figure S8. Mycoplasma detection in cell lines used in this study.** All cell lines used in this study have been tested for the presence of Mycoplasma using MycoStrip™ from Invivogen. The respective cell lines are labelled on and below each test strip. One red line indicates positive for internal control and negative for sample tested. Double red line (positive control) indicates positive for mycoplasma. For hNK cells, Primocin® and Plasmocin® Prophylactic were added to every fresh batch of isolated hNK cells.

**Supplementary Table S1. Primers used in this study for RT-qPCR analyses.**

| Gene | Primer | Sequence (5’-3’) | Tm (°C) |
|---|---|---|---|
| GAPDH | Forward | **ATG TTT GTG ATG GGT GTG AA** | 60.0 |
| | Reverse | **ATG CCA AAG TTG TCA TGG AT** | 60.0 |
| c-Myc | Forward | **CCT GGT GCT CCA TGA GGA GAC** | 60.0 |
| | Reverse | **CAG ACT CTG ACC TTT TGC CAG** | 60.0 |
| WWTR1 | Forward | **CCT GGT GCT CCA TGA GGA GAC** | 60.0 |
| | Reverse | **ATT CGA ATG CGC CAA GAG** | 60.0 |
| YAP1 | Forward | **TGT CCC AGA TGA ACG TCA CAG C** | 58.0 |
| | Reverse | **TGG TGG CTG TTT CAC TGG AGC A** | 58.0 |

**Supplementary Table S2. List of antibodies used in this study.**

| Antibody | Catalogue No. | Manufacturer | RRID[@] |
|---|---|---|---|
| CD3-eFluor450 | 48-0032-82 | eBioscience | RRID: AB_1272193 |
| CD56-APC | 17-0567-41 | eBioscience | RRID: AB_10596498 |
| CD107a-eFluro610 | 61-1079-42 | eBioscience | RRID: AB_2574572 |
| Ki67-PE | 12-5699-42 | eBioscience | RRID: AB_10688373 |
| PD1-PE | 130-120-382 | Miltenyi Biotec | RRID: AB_2752069 |
| TIGIT-PE | 130-116-814 | Miltenyi Biotec | RRID: AB_2751336 |
| NKG2A-APC | 130-113-563 | Miltenyi Biotec | RRID: AB_2726170 |
| NKG2D-PE | 130-123-709 | Miltenyi Biotec | RRID: AB_2819516 |
| TLR3-APC | 130-110-383 | Miltenyi Biotec | RRID: AB_2657871 |
| Perforin-eFluro 450 | 48−9994−42 | Invitrogen | RRID: AB_2574145 |
| Granzyme B-APC | 372204 | BioLegend | RRID: AB_2687028 |
| KIR3DL1-APC | 312716 | BioLegend | RRID: AB_2563360 |
| KIR2DL1-PE | 374903 | BioLegend | RRID: AB_2832735 |
| CD16-PE | 302007 | BioLegend | RRID: AB_314207 |
| TAZ | 83669 | CST | RRID: AB_2800026 |
| LATS-1 | 9153 | CST | RRID: AB_2296754 |
| β-actin | 3700 | CST | RRID: AB_2242334 |
| p-TAZ (S89) | 59971 | CST | RRID: AB_2799578 |
| p-LATS-1 (S909) | 9157 | CST | RRID: AB_2133515 |
| YAP | 4912 | CST | RRID: AB_2218911 |
| p-YAP(S127) | 4911 | CST | RRID: AB_2218913 |
| p-MLC2 | 3674 | CST | RRID: AB_2147464 |
| p-Erk1/2 | 9101 | CST | RRID: AB_331646 |
| Myosin Light Chain | 3672 | CST | RRID: AB_10692513 |
| Erk1/2 | CST | CST | RRID: AB_330744 |
| Anti-RhoA | ARH04 | Cytoskeleton, Inc | RRID: AB_2728698 |

[@] RRID, Resource Reference ID
[@] CST, Cell Signaling Technology
