## Supplementary Figures 1-8 for "Natural Killer cell contractility and cytotoxicity is driven by TLR3 agonist-mediated TAZ cytoplasmic sequestration"

Figure S1. NK cells express TAZ but not YAP

A (TAZ/ WWTR1)

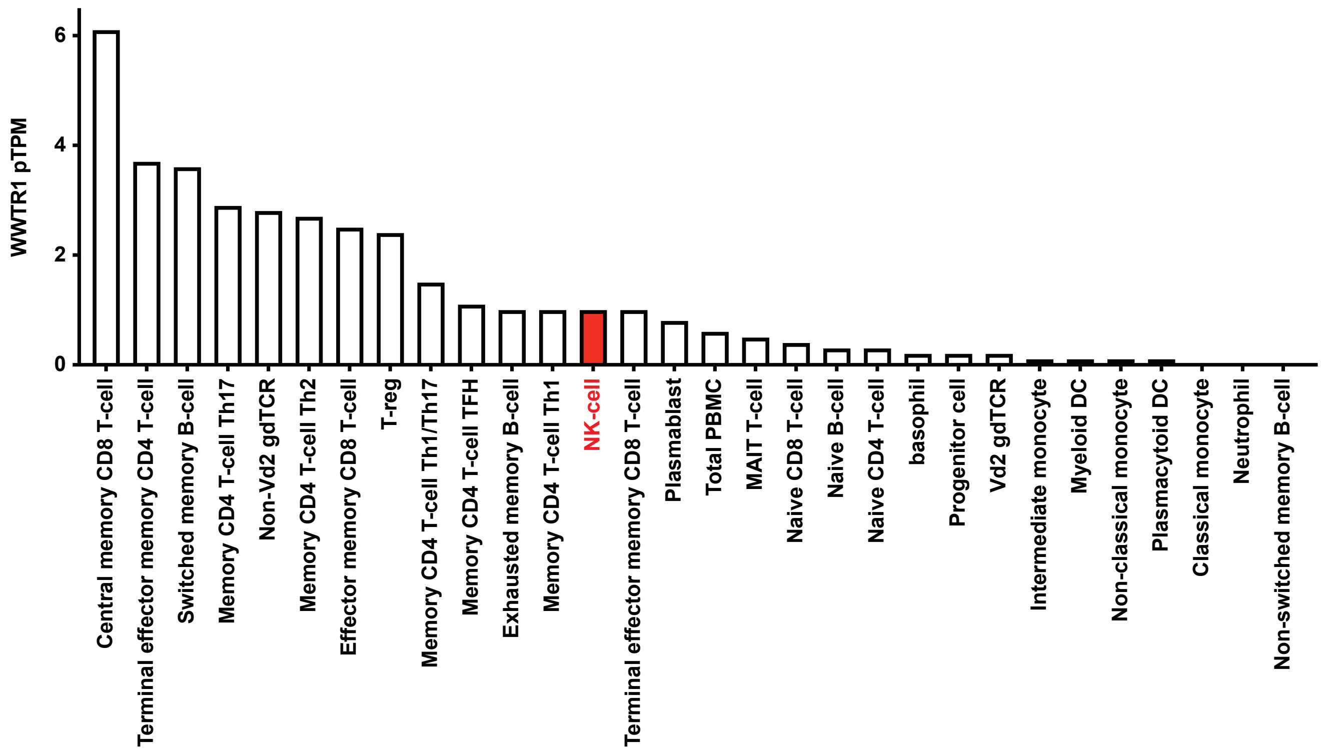


B (YAP)

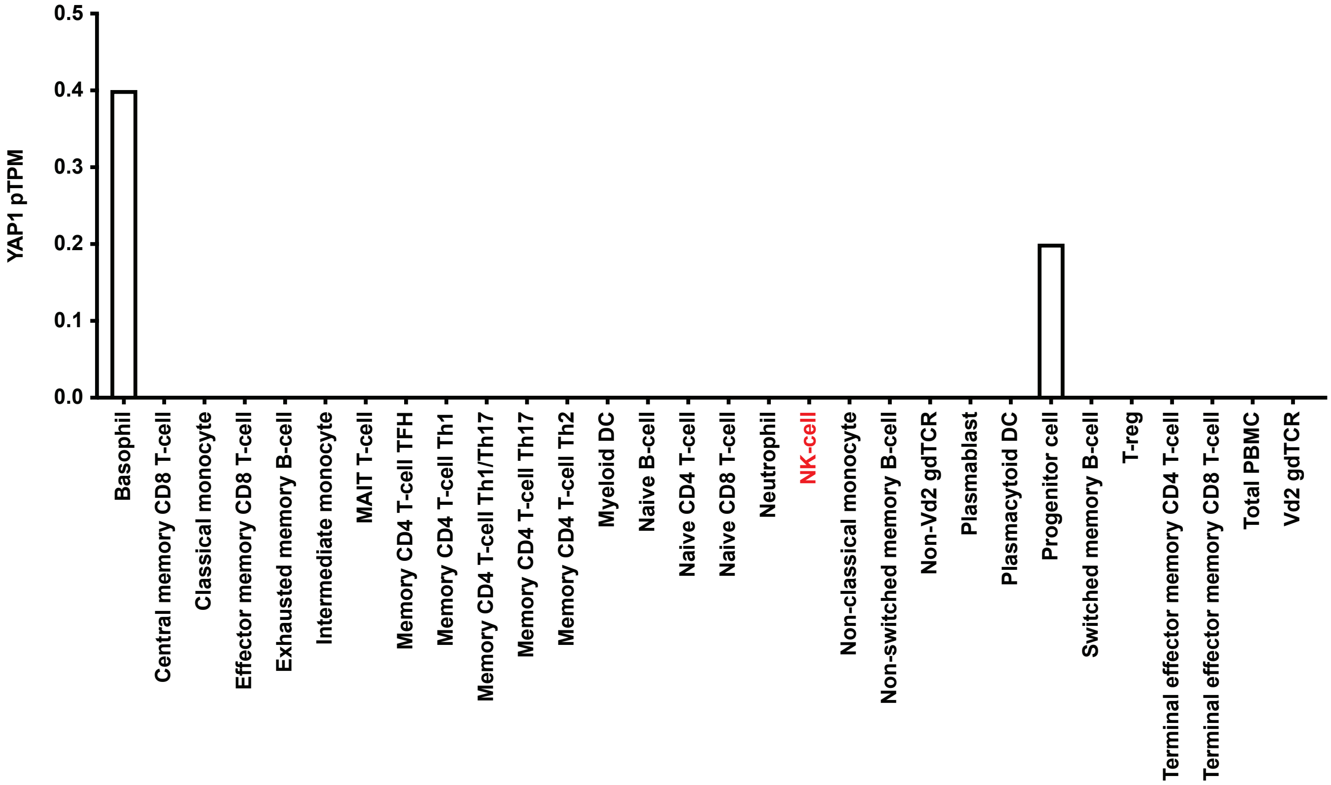

**Figure S2. Gating strategy for FVD506 staining of dead cells**

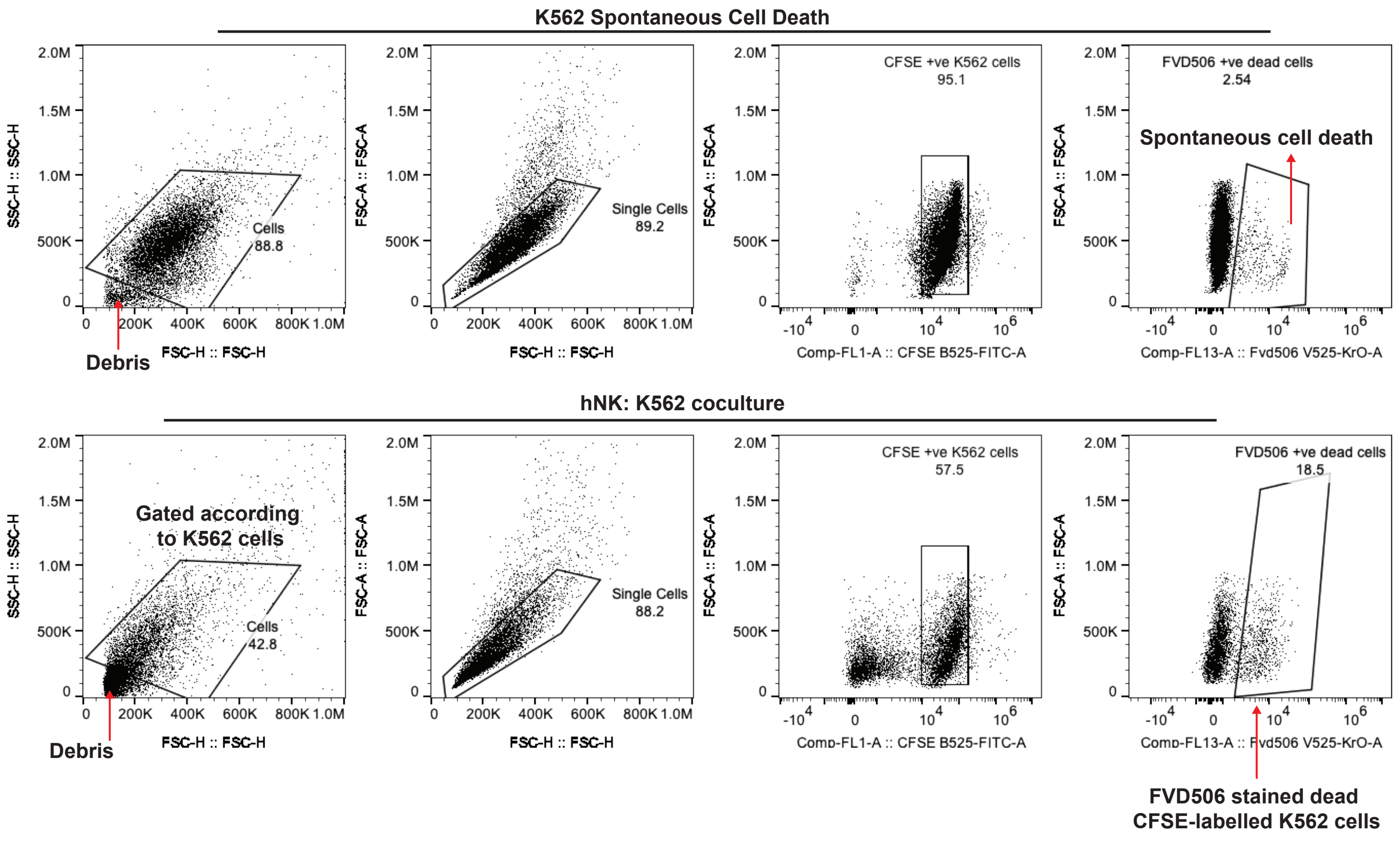

**Figure S3. Hiltonol increases TLR3 expression, induces Eomes nuclear localization in NK cells, and promotes NK:K562 binding**

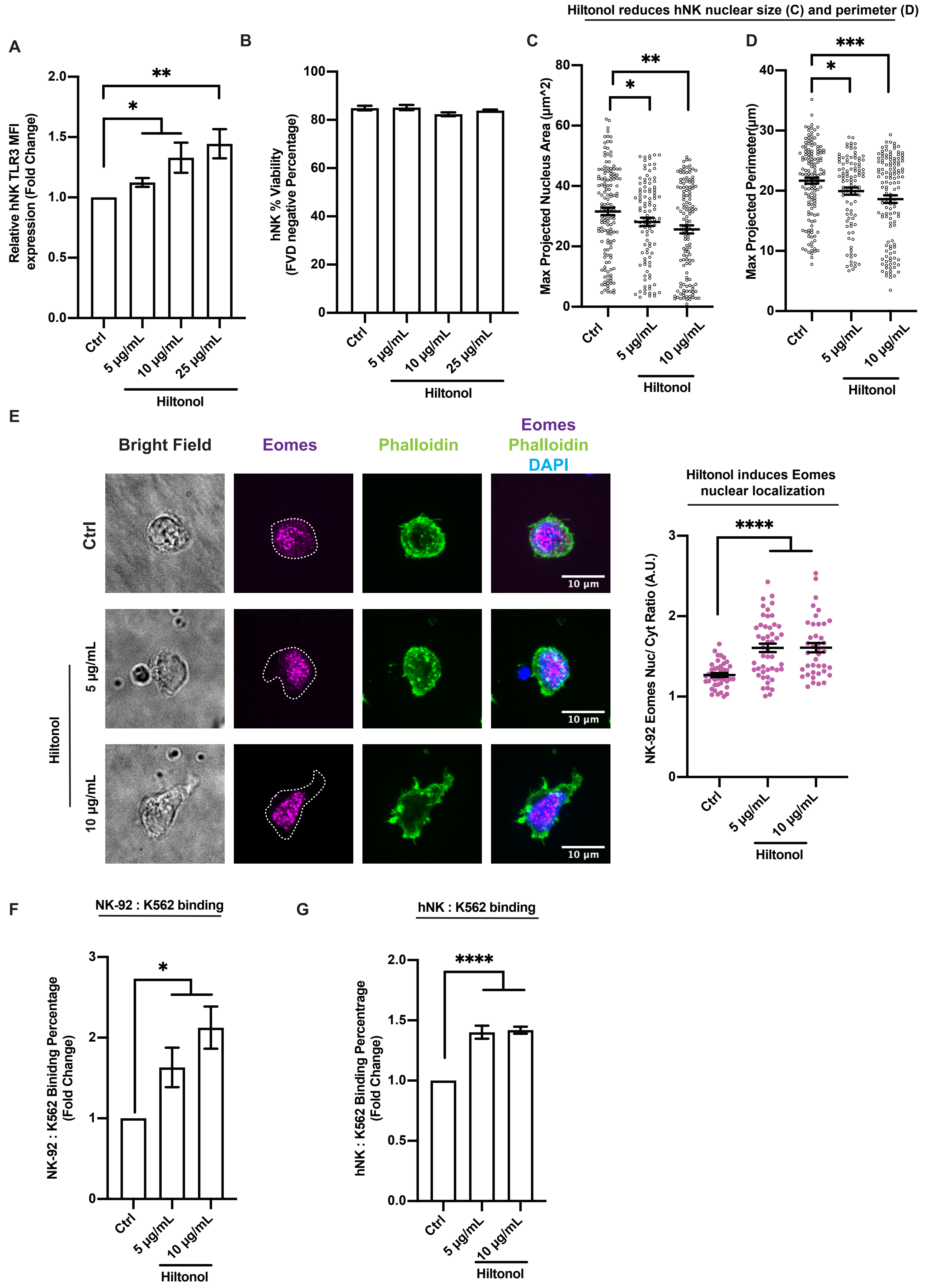

### Figure S4. Hiltonol and G3798 treatment do not alter hNK surface CD56 and CD16 expressions

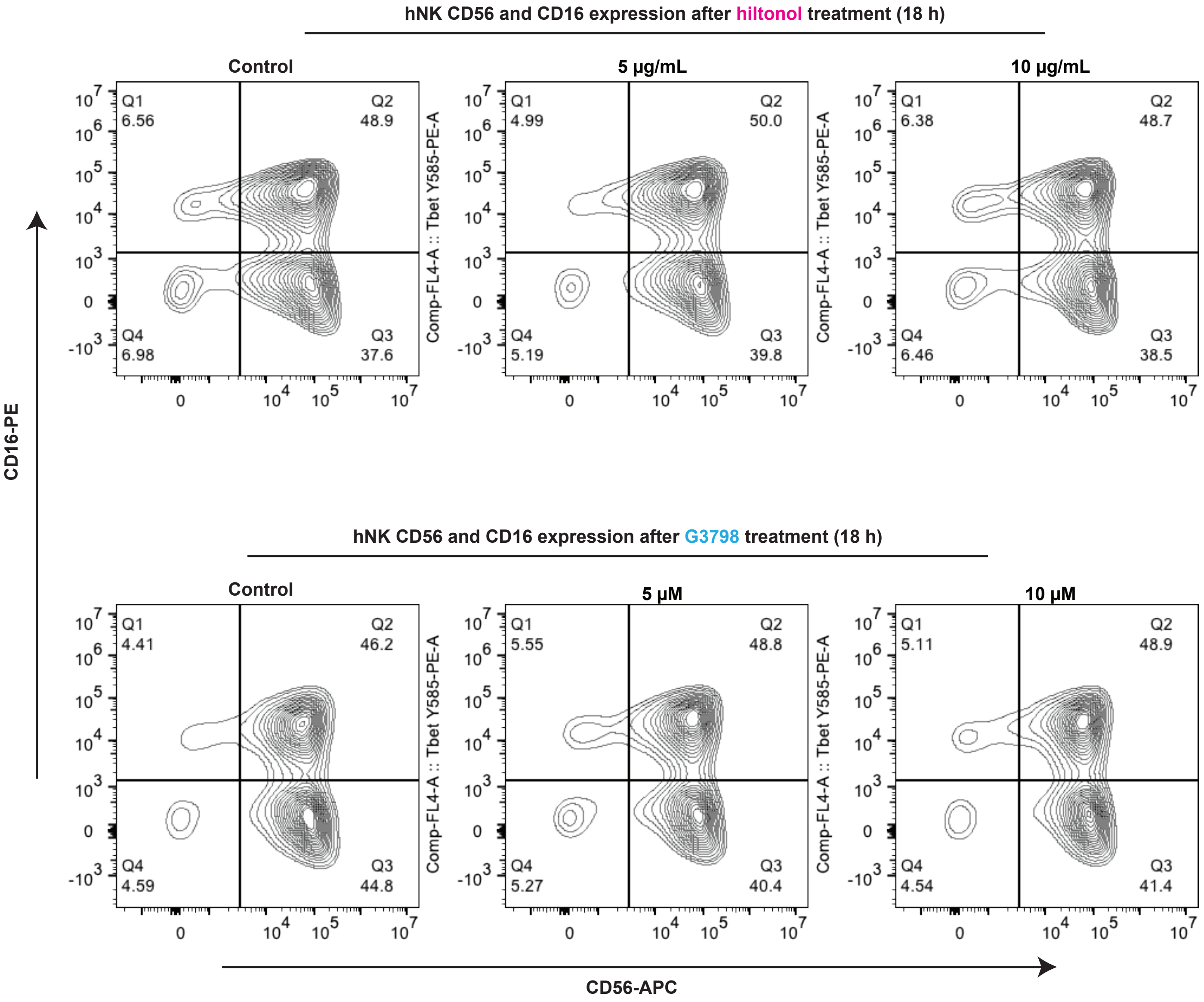

**Figure S5. High dose of c-Myc inhibitor (G3798) induces cell death on less metastatic cancer cells**

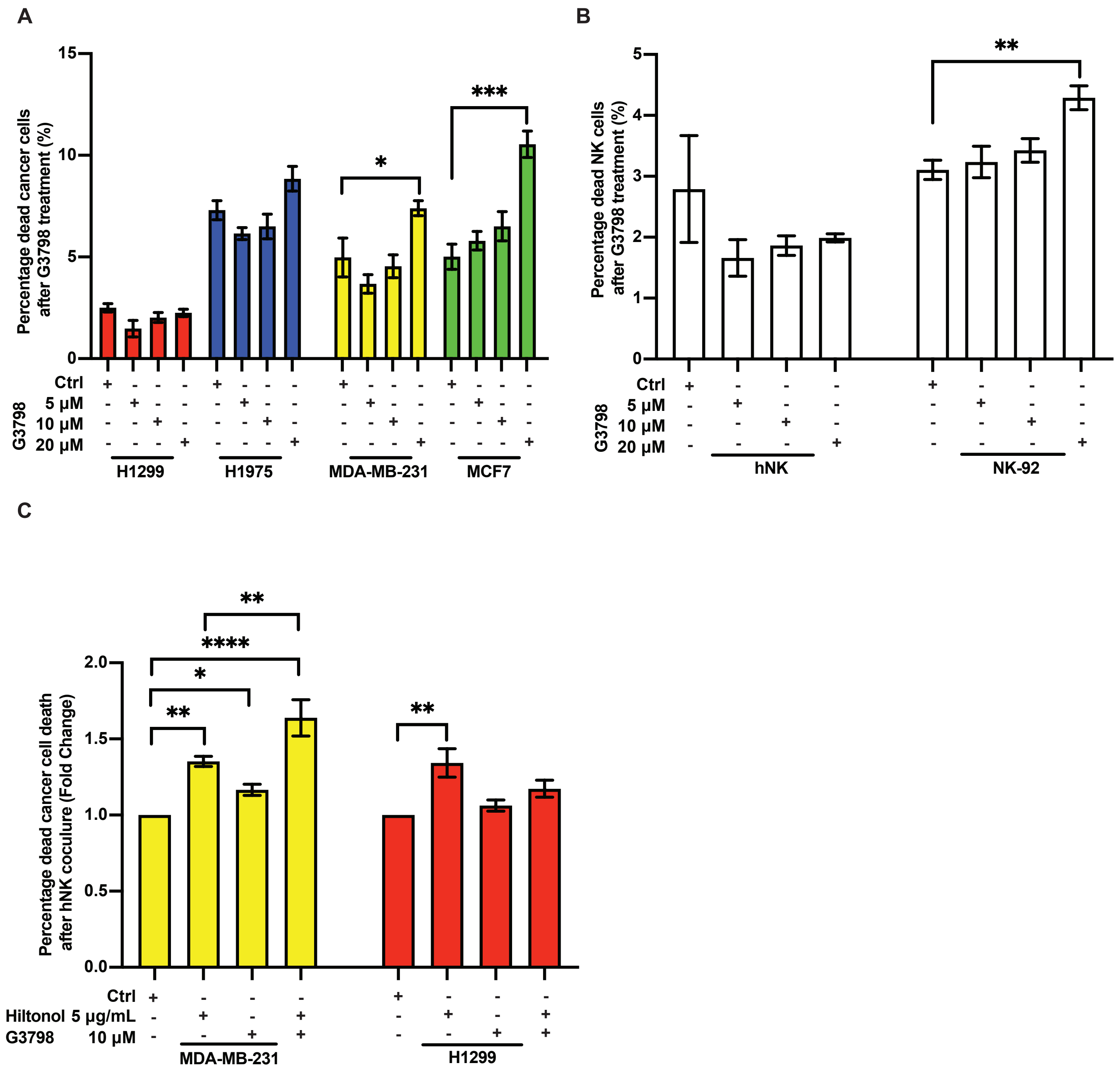

**Figure S6. Representative gating strategy for flow cytometry experiments used in the study**

**A**

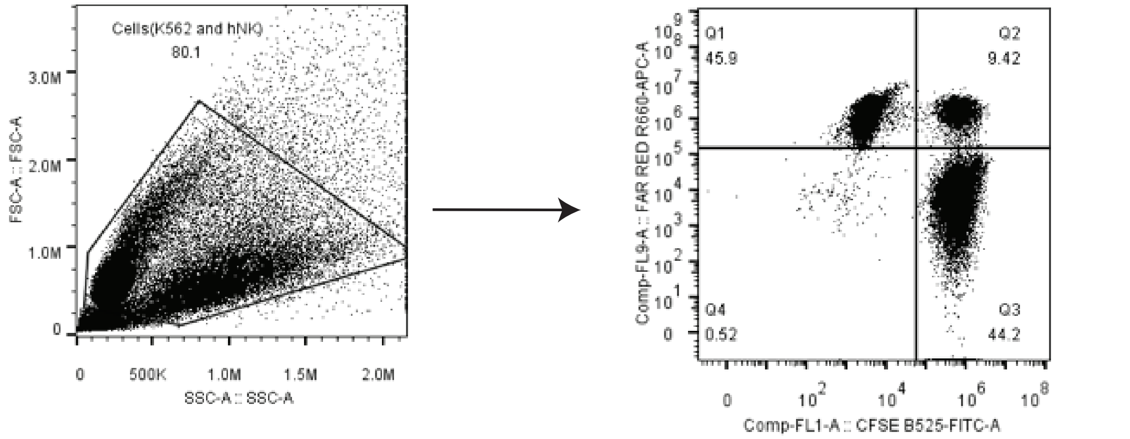


**Q1: K562-Far red APC Cells**
**Q2: K562-Far red APC: hNK CFSE-Cells**
**Q3: hNK-CFSE Cells**

**B**

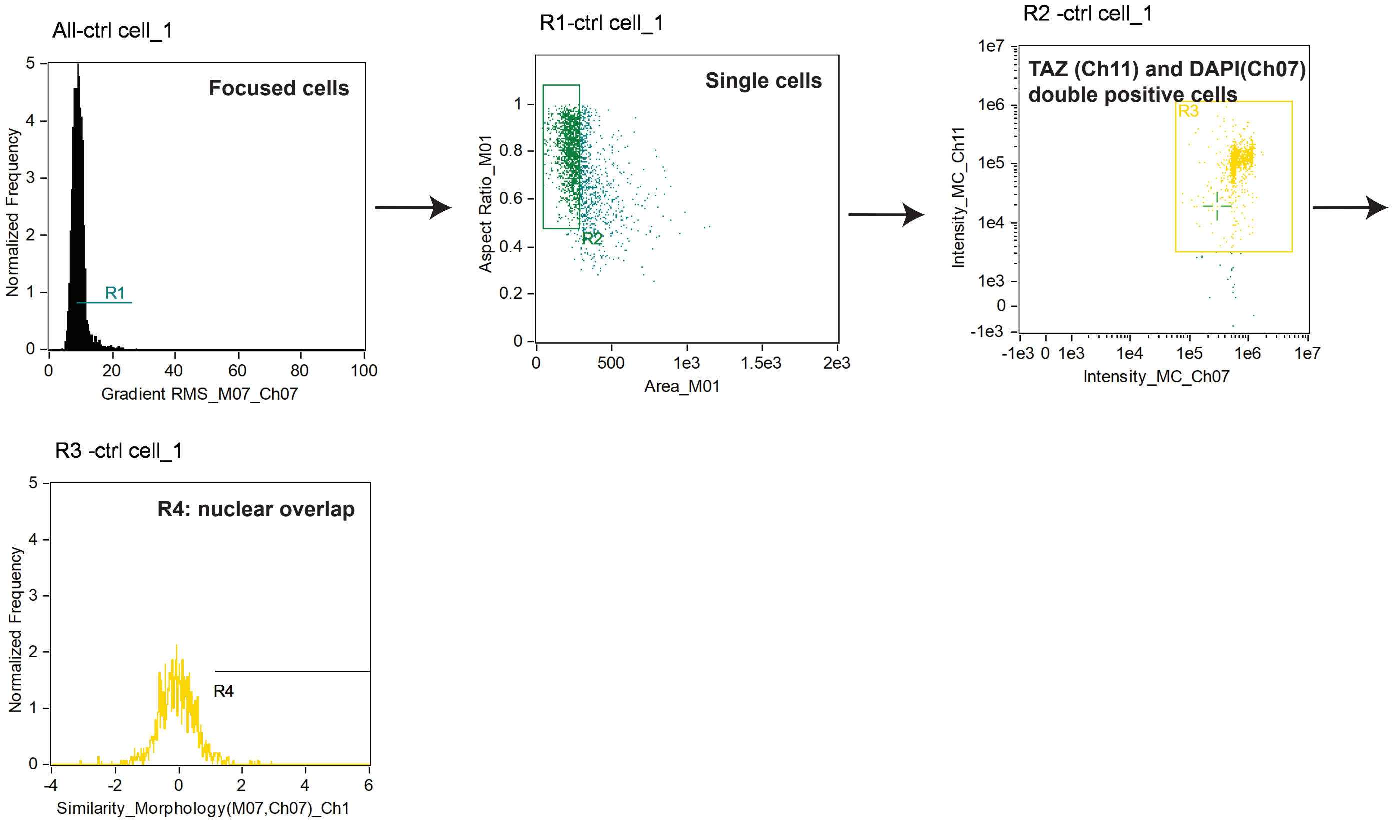

**Figure S7. Flow cytometry antibody controls (Fluorescence minus one, FMO) used in this study**

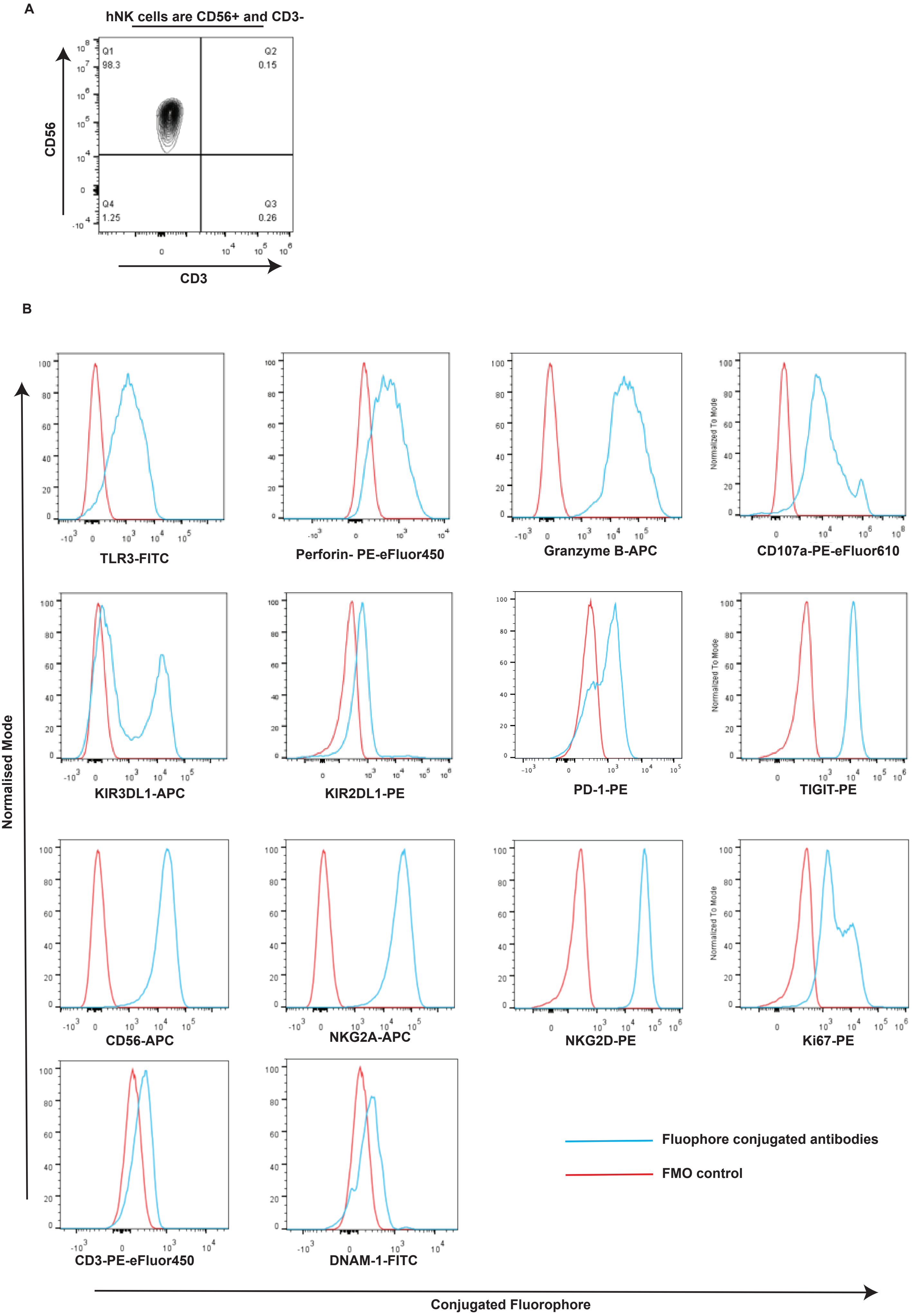

**Figure S8. Mycoplasma detection in cell lines used in this study**

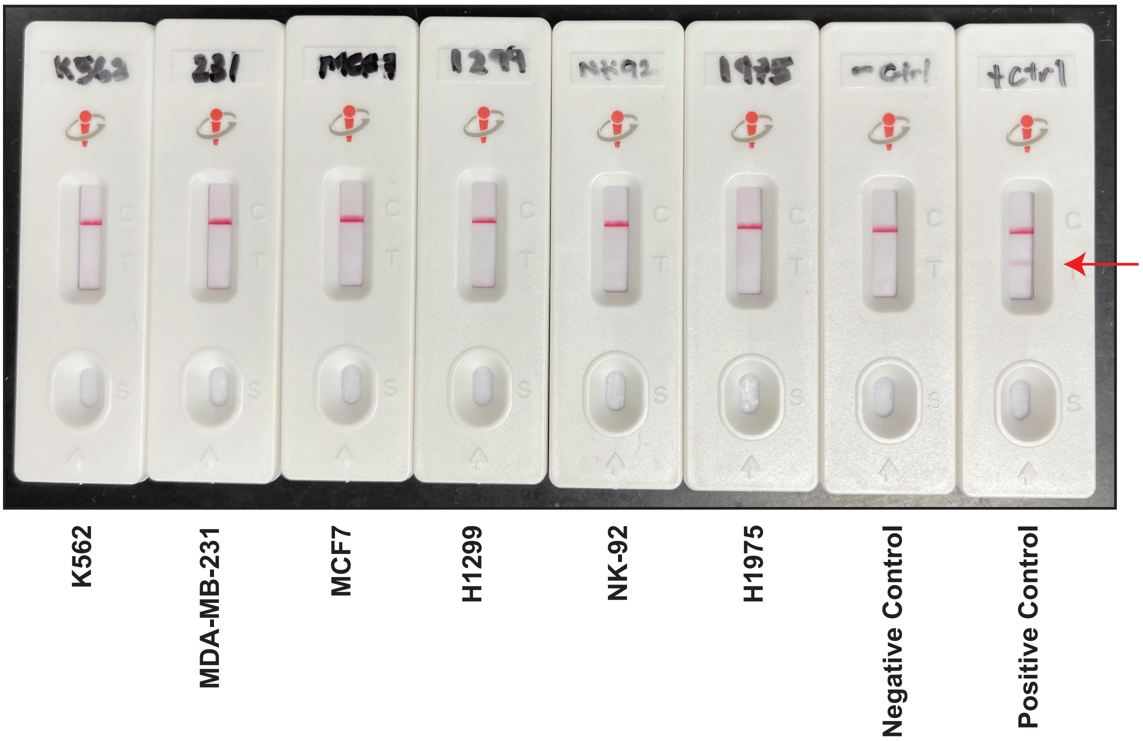
